# Learning fragment-based segmentation of binding sites from molecular dynamics: a proof-of-concept on cardiac myosin

**DOI:** 10.64898/2026.02.13.703009

**Authors:** Yu-Yuan Yang, Richard W. Pickersgill, Arianna Fornili

## Abstract

The geometric and chemical features of protein binding sites tend to change as a consequence of conformational dynamics. In the ligand-unbound (apo) state, a binding site might be only transiently organised in a way that can accommodate a given ligand, with the relevant regions of the protein coming together in a suitable arrangement only in a subset of conformations. Ligand binding itself can also induce further changes in the binding site.

Because most ligands can be decomposed into smaller fragments, we hypothesised that mapping onto the binding site surface the propensity of binding specific fragments could be used to monitor changes in the overall ability of the site to bind a ligand. This task can be formulated as semantic segmentation, which can now be performed successfully using deep learning methods.

Here we introduce the Fragment-Based protein Ensemble semantic Segmentation Tool for Myosin (FragBEST-Myo), a deep learning method based on a 3D U-Net architecture, trained to partition the omecamtiv mecarbil (OM) binding site of cardiac myosin into fragment-specific regions using only local shape and physico-chemical features.

The model was trained on labelled Molecular Dynamics (MD) trajectories of OM-bound myosin in both post-rigor (PR) and pre-power-stroke (PPS) states, achieving an accuracy of ~95% and a mean Intersection over Union (mIoU) > 0.75 on unseen trajectories from both states. When applied to apo trajectories, FragBEST-Myo-derived descriptors produced rankings consistent with similarity to holo conformations. Moreover, selecting apo frames based on FragBEST-Myo ranking increased the chance of recovering holo-like OM docking poses relative to randomly chosen control frames, supporting its use as a screening tool for ensemble docking. Beyond frame selection, fragment maps provide a compact representation to assess docking poses and to guide fragment-based design.

Our proof-of-concept provides a basis for developing future general models applicable to a broader range of proteins and ligands, with the fragment-based formulation offering a natural route to generalisation.

**Graphical Abstract:** 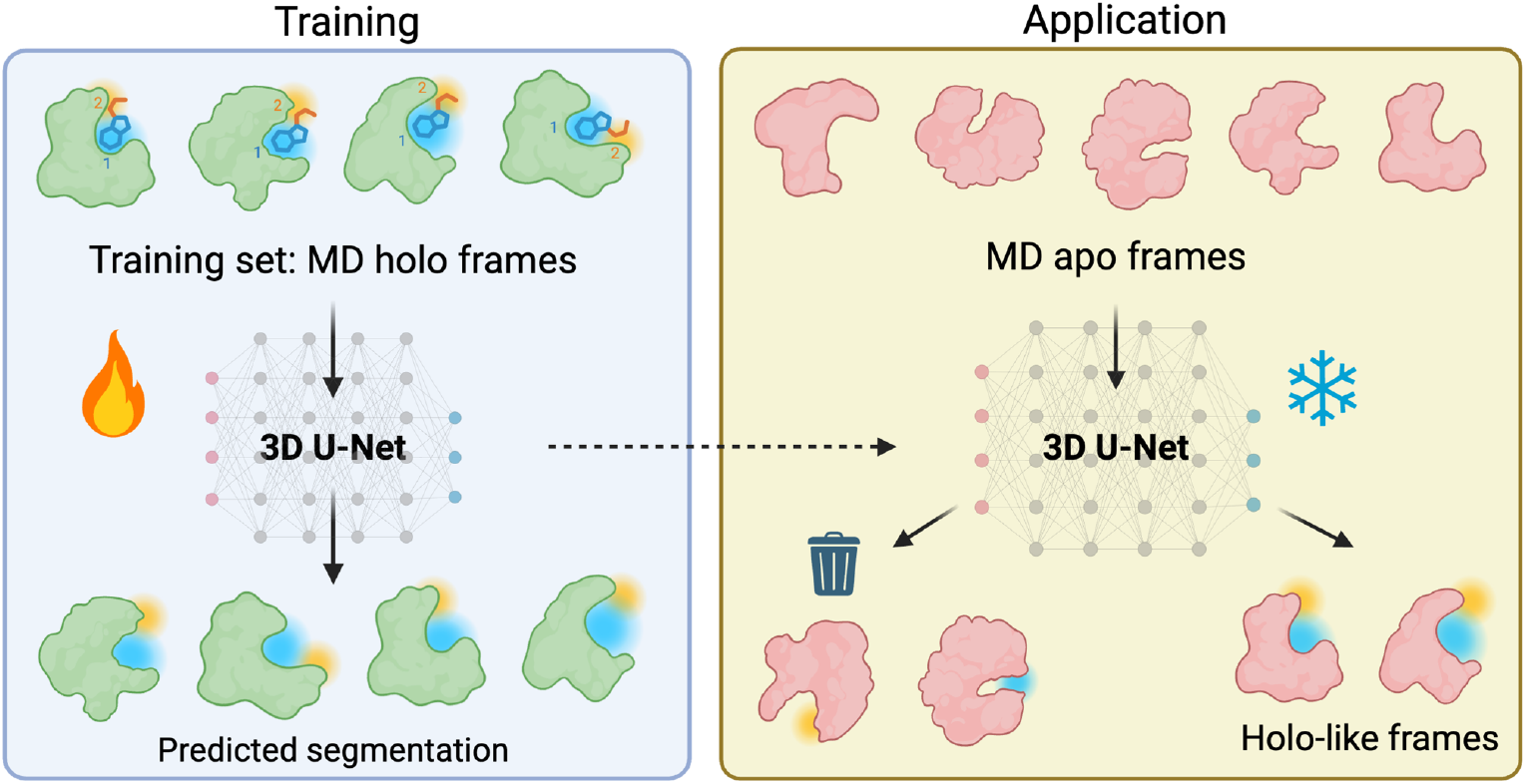

## Introduction

Protein binding sites are not static structures but dynamic regions whose ability to accommodate a ligand depends on complementarity of local shape and physico-chemical properties. Both the geometry of the pocket and the distribution of chemical features can change as the protein explores its conformational space. In the ligand-unbound (apo) state, the binding site might be only transiently organised in a way that can accommodate a given ligand, with the relevant regions of the protein coming together in a suitable arrangement only in a subset of conformations (conformational selection)^1,2^. Ligand binding itself can also induce changes in the binding site (induced fit) and different ligands might even promote different ligand-bound (holo) structures.

Since most ligands can be viewed as combinations of smaller fragments, it would be natural to describe the binding site in terms of regions that can bind specific fragment types. Mapping the propensity of binding specific fragments onto the binding site surface could be used to monitor changes in the overall ability of the site to bind a given ligand. Conversely, fragment-binding maps could guide the design of new ligands optimised to bind specific conformations.

In this work, we investigated whether deep learning (DL) models for semantic segmentation can be trained to partition a binding-site surface into regions associated with defined ligand fragments, using only the local protein environment (shape and physico-chemical properties) as input, and taking into account structural variations observed in molecular dynamics (MD) simulations. We adopted a 3D U-Net model^3,4^, a widely used architecture for image segmentation tasks in medical imaging^5,6^, which has also been successfully applied in the context of binding-site detection in proteins^7–9^.

At this stage, our goal was not to develop a general approach across many targets, but to address more focused questions on a single, well-characterised system. Can a model trained on holo trajectories recover an appropriate fragment-based segmentation of the binding site across the conformational landscape sampled by the apo protein? Does the predicted segmentation contain enough information to distinguish frames that are compatible with ligand binding from those that are not, such that it can be used to guide the selection of frames for structure-based modelling of protein-ligand interactions? This second aspect is important because the success of many structure-based drug discovery approaches, such as ensemble docking-based virtual screening^10,11^, relies on including structures that can bind the ligand of interest^12,13^. MD simulations, whether unbiased or enhanced^11,13–15^, can sample such conformations even when started from apo structures, but identifying them among the generated frames remains challenging.

To answer these questions, we considered cardiac myosin bound to the first-in-class cardiac selective activator omecamtiv mecarbil (OM)^16,17^. Myosin is known to adopt different conformational states during its functional cycle, including the post-rigour (PR) and pre-power stroke (PPS) states, with OM binding to a key region of the motor domain at the interface between different subdomains (Figure 1A). We developed a model, the Fragment-Based protein Ensemble semantic Segmentation Tool for Myosin (FragBEST-Myo), trained on a labelled dataset composed of multiple MD simulations of OM-bound cardiac myosin in both PR and PPS states. For each MD frame, the OM binding-site surface was computed, voxelised, and each voxel labelled according to the closest OM fragment. FragBEST-Myo was first trained to predict these labels and tested against unseen holo trajectories. Subsequently, it was applied to apo trajectories sampling the individual states (PR or PPS), and a steered MD (SMD) trajectory in which the protein was driven from PR to PPS, sampling multiple states within a single simulation. The predicted binding site segmentation was finally used to identify apo frames likely to bind OM, which were then tested with molecular docking calculations.

**Figure 1.**
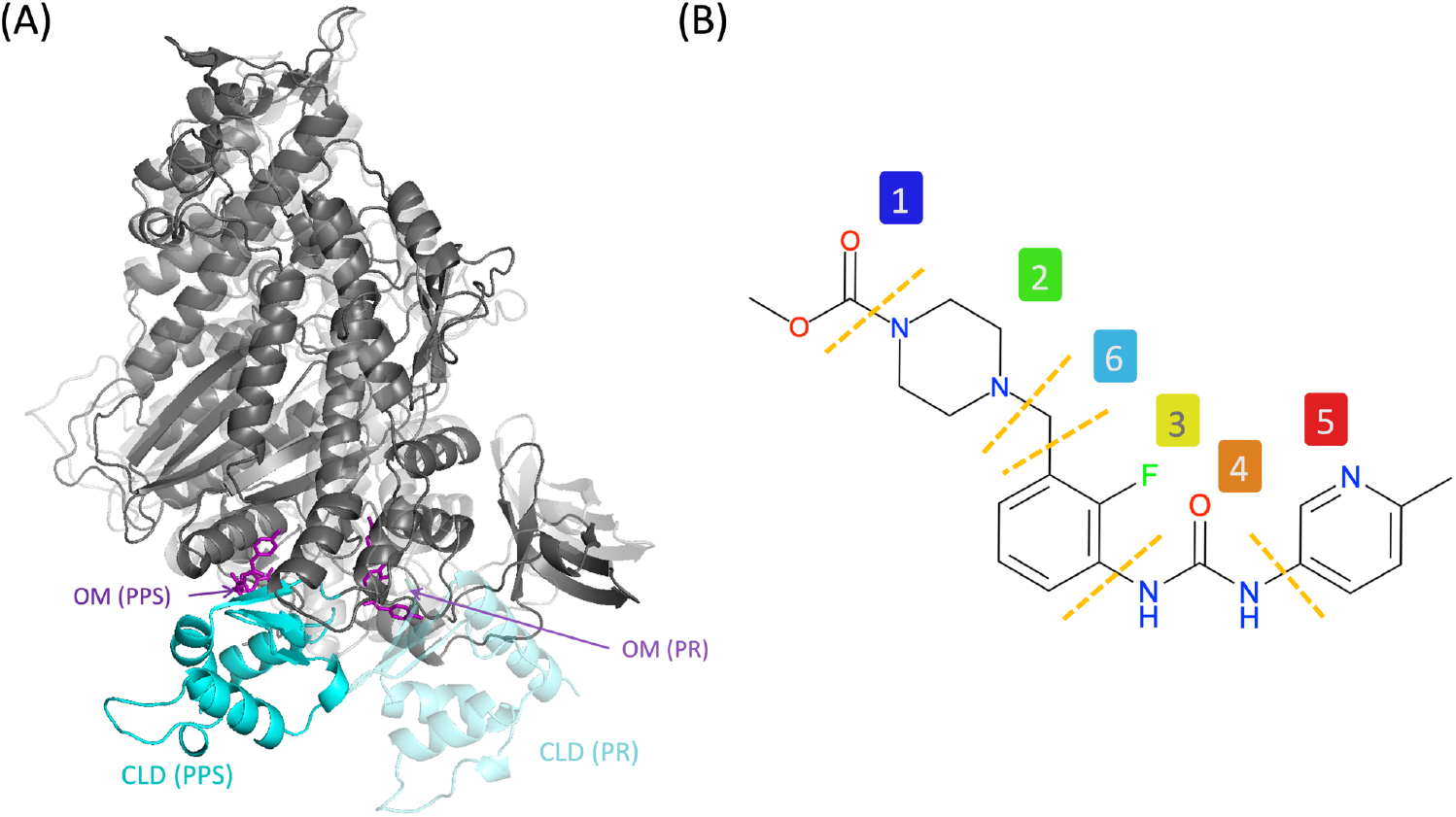
Cardiac myosin and OM. (A) Cartoon representation of cardiac myosin in the PPS (opaque) and PR (transparent) conformational states (starting structure of MD simulations^16,17^). The converter subdomain and the N-terminal portion of the lever arm (CLD) are highlighted in cyan, while OM is represented as purple licorice. (B) Decomposition of OM into C fragments.

## Methods

### Dataset preparation

The datasets used for training, validation, and testing of the DL models were built using previously published MD simulations of the motor domain of cardiac myosin (CM) in the post-rigour (PR)^17^ and pre-power stroke (PPS)^16^ states (Table S1). A total of 4 OM-bound (holo) and 4 OM-unbound (apo) CM replicas were used for each state. Frames were extracted every 100 ps from trajectories of about 300-ns (with slight variations in total length across systems). In addition, the final model was tested on a 200-ns Steered MD (SMD) simulation of the PR-to-PPS transition of apo CM^18^ sampled every 100 ps.

All the PPS, PR, and PR-to-PPS trajectories were first aligned to a single structure (holo PPS energy-minimised structure) based on the C_α_ atoms of residues 1 to 700. The converter and lever arm subdomains of the protein were not used for the alignment as they adopt significantly different orientations in the two conformational states^18^ (Figure 1A). Each trajectory was then aligned to its own first frame using only the residues in the OM pocket (the initial alignment ensured that the first frames of all the trajectories were globally aligned). The OM pocket was defined as the set of residues found within 5 Å of the ligand for at least 50% of the frames for holo simulations, and the set of residues within 5 Å of OM in the energy-minimised holo structure of the corresponding state for apo simulations (Table S1). The geometric centre of all the pocket atoms was used as the pocket centre. MDAnalysis^19^ was utilised to process the MD trajectories.

For each frame, the region of interest (ROI) was defined as a 32-Å-diameter sphere positioned at the pocket centre. A mesh representation of the protein surface within the ROI was generated using the MaSIF^20,21^ framework, where the solvent-excluded surface (SES) is calculated using MSMS^22^ and discretised into a mesh with 1-Å spacing. For each vertex, three physico-chemical features were calculated as in MaSIF^20,21^, measuring Poisson–Boltzmann continuum electrostatics, hydrogen bond potential, and hydrophobicity. Vertices and features were written into files using PyMesh^23^.

For holo frames, the ROI vertices were also assigned a label according to their relative position to the ligand. OM was first manually decomposed into six different fragments using RDKit^24^, and a class *c* was created for each fragment (Figure 1B). Every vertex located within a 5 Å radius from the ligand’s heavy atoms was assigned a label corresponding to the class of the nearest fragment (*c* = 1-6). All other vertices within the ROI were categorised as background (*c* = 0), leading to a total of 7 classes.

Before model training, the vertices in the ROI were voxelized into a 64 × 64 × 64 3D grid spanning a 32 × 32 × 32 Å^3^ region. The grid spacing was set to 0.5 Å to minimise the loss of vertices while keeping the computation efficient. Each ROI voxel was assigned the same label as the corresponding ROI vertex. Voxels outside the ROI were assigned the background class label. Each of the three vertex features was Z-score normalised and assigned to a separate channel (0–2). A fourth channel (channel 3) was added to store the occupancy of the voxels (set to 1 for voxels with ROI vertices and 0 otherwise). This channel was used both for training and validation as a mask for loss calculation and metric evaluation.

### Deep-learning model training

3D U-Net convolutional networks^3,4^ are widely used for 3D biomedical image segmentation tasks^5,6^. They have also been adopted for voxel-based representations of binding sites in prediction tasks^9^. Here, we used a PyTorch implementation^25^ of a 3D U-Net. Four channels (one for each vertex feature type plus the occupancy mask) were used for the input and seven (one for each class of vertex labels) for the output, giving tensor shapes of 4 × 64 × 64 × 64 and 7 × 64 × 64 × 64, respectively. Compared to the original implementation^4^, exponential linear units (ELUs)^26^ replaced rectified linear units (ReLUs) as activation function in convolutional building blocks to speed up learning and a softmax function was used as final activation. The optimisation process was carried out using the Adam optimiser. GPU memory constraints required the adoption of gradient accumulation techniques to achieve a batch size of 4 or 8 during training^27^. The learning rate was established at 0.0001, with the training extending to a maximum of 600 epochs. To address class imbalance, predefined weights ([0.25, 1, 1, 1, 1, 1, 1]) were assigned to each class in the calculation of the loss function. Furthermore, a combination of generalised dice loss^28^ and focal loss^29^ (GDF loss) was used (with focusing parameter γ = 2 and trade-off weight λ = 1 for both losses). Considering the vertices were sparsely mapped to the voxel grid, a masked version of GDF loss, where the occupancy channel served as the mask, was implemented using MONAI^30^ to accelerate model training (Figure S1 and Supplementary Methods).

Model performance was assessed with two commonly used segmentation metrics, accuracy and intersection over union (IoU)^5,31^. The mean IoU (mIoU), calculated as average IoU across classes, was utilised for multi-class segmentation evaluation. K-fold cross-validation was used with K = 3, 5, 6, or 10. 3D random rotation data augmentation was also implemented during training using MONAI^30^. For a given K value and training setting, the maximum mIoU values observed during training were recorded for each fold when calculated on the training (best train mIoU) and validation (best validation mIoU) sets. Training and validation performance were reported as average and standard deviation of the corresponding best mIoU values across the K folds. The model with the highest validation mIoU across the three folds was then selected for testing. Testing performance was reported as average mIoU across testing set frames together with the standard deviation.

Frames in the training set were randomly rotated around the x, y, or z axis with a probability of 0.5, using rotation angles sampled from one of four ranges: 0° (no data augmentation applied), −45° to 45°, −90° to 90°, or −180° to 180°. PyTorch Ignite^32^ was used to manage the training process, including multi-GPU distributed computing, model saving, and performance evaluation after each epoch. Fixed random seeds were used for all stochastic operations (e.g. dataset sampling and splitting, random rotations, initialisation of model parameters when trained from scratch) to ensure reproducibility.

Models were initially trained using only the PPS training set. Once the hyperparameters were optimised (including the choice of loss functions, the training set size, the number of folds in K-fold cross-validation and the rotation range for data augmentation), a more generalised model was trained on both PPS and PR training sets, starting from the pretrained weights of the best PPS model.

### Descriptors and scoring function

A scoring function was designed to rank trajectory frames based on their binding site segmentation predicted by the best DL model. The overall score of each frame was calculated using four descriptors (Table 1): the number of distinct labels assigned by the model to all the ROI vertices in the frame (D_1_), the fraction of vertices labelled with any non-background class (D_2_), the fraction of vertices labelled with a specific non-background class, averaged over all non-background classes counted in D_1_ (D_3_), and a volume-based descriptor, the averaged capped HoloSpace score (D_4_).

**Table 1.** Summary of D1-4 descriptors with typical ranges observed in this study.

| Descriptor | Meaning | Range |
| --- | --- | --- |
| D<sub>1</sub>: Number of distinct labels | Number of different fragments predicted to bind to the binding site. | 1 (only one class detected, usually background) to 7 (all classes detected) |
| D<sub>2</sub>: Fraction of non-background vertices | A higher value means that a higher proportion of the ROI vertices is predicted to be involved in the binding of the ligand. | 0% (no vertex with a label $\neq 0$) to ~30% (total fraction of non-background vertices in a holo conformation) |
| D<sub>3</sub>: Average fraction of vertices with a specific non-background label | A higher value means that on average a higher proportion of a segmented region is | 0% (no vertex with a label $\neq 0$) to ~5% (average of individual fractions of non- |
|  | predicted to be involved in the binding of its corresponding fragment. | background vetices in a holo conformation) |
| D4: Averaged capped HoloSpace score | A higher score indicates that the segmented regions are sufficiently large to accommodate their corresponding fragments. | 0 (not enough space to accommodate any fragment) to 1 (enough space for all fragments) |

The following equations were used for D_2_ and D_3_:

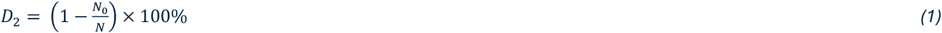

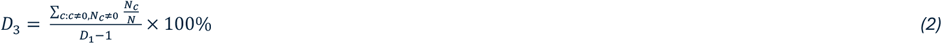

where *N*_*c*_ is the total number of vertices labelled with class *c* (e.g., *N*_0_ is the total number of ROI vertices labelled as background) and *N* is the total number of ROI vertices in the frame.

Inspired by the pocket-fragment complementarity scoring in the Protein-Protein Interaction interface mapping tool AlphaSpace^33^, the HoloSpace concept was introduced to assess whether a region delimited by a group of vertices *V*_*c*_ labelled with a non-background class *c* was large enough to accommodate the corresponding fragment. The raw HoloSpace (*rHS*_*c*_) was defined as the intersection between the solvent-accessible space and a sphere placed at the geometric centre (*g*_*c*_) of the *V*_*c*_ vertices, with a radius equal to the maximum distance from *g*_*c*_ to any of the *V*_*c*_ vertices (Figure 2A). To take into account that adjacent raw HoloSpaces may overlap, a corrected HoloSpace volume *cHSV*_*c*_ was calculated (Figure 2B). For a given *rHS*_*c*_, the volume of the intersection with every other *rHS*_*p*_ (with p≠c) was first determined. The top two intersecting volumes were then halved and subtracted from the *rHS*_*c*_ volume, yielding *cHSV*_*c*_. This procedure assumes that substantial overlap arises from at most two adjacent regions, which is a reasonable assumption considering the linearity of the OM chemical structure.

**Figure 2.**
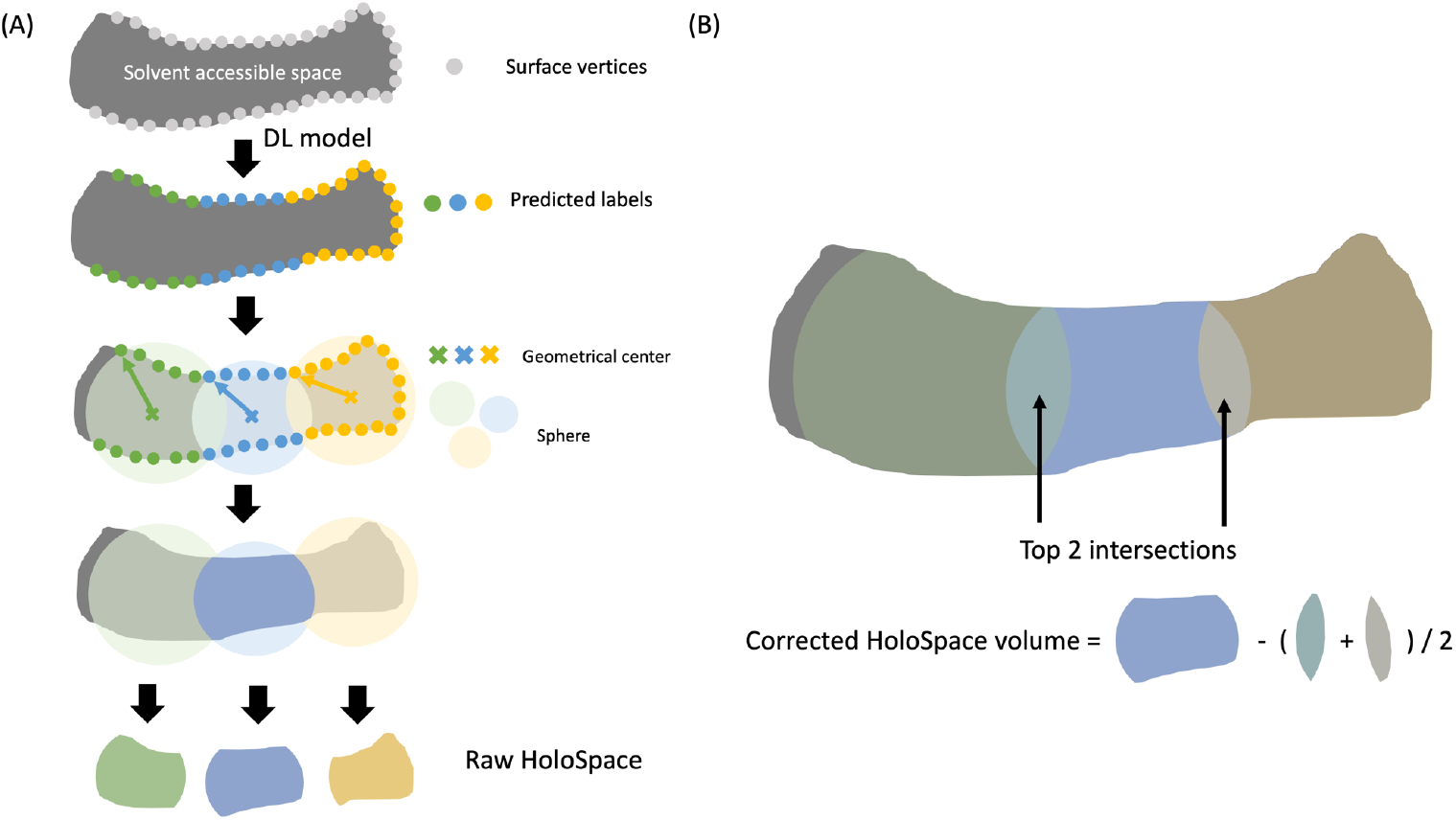
Schematic representation of Raw (A) and Corrected (B) HoloSpace volumes. The solvent-accessible space is delineated by the ROI vertices.

The corrected HoloSpace volume *cHSV*_c_ was then compared with the reference volume of the corresponding fragment determined by the RDKit^24^ (*refV*_*c*_) to calculate the capped HoloSpace score *cHSS*_*c*_:

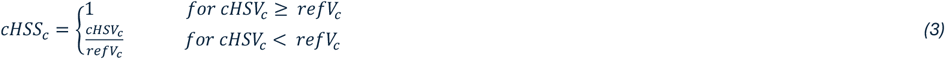

Finally, the averaged capped HoloSpace score (D_4_) was calculated as the average *cHSS*_c_ over the non-background classes.

Each descriptor was z-score normalised, and an overall score was obtained for each frame by summing the normalised scores from all descriptors with equal weighting. Frames were then assigned a rank R_holo_ based on the overall score. Before ranking, a filter was applied to remove frames with anomalies in the mesh generation or overestimated HoloSpace volumes as described in the supplementary information (“Pre-ranking filters”).

### Ensemble docking

The performance of the R_holo_ ranking was assessed using molecular docking calculations. A set of structures (top100 sets) was built by selecting the 100 top-ranking frames from the apo PPS or PR trajectories. As a control, 100 frames were randomly sampled from the same trajectories (sample100 sets). For each selected frame, pairwise Root Mean Square Deviation (RMSD) values were calculated between that frame and all holo PR or PPS structures used for training (RMSD_holo,X_, with X=PR or PPS). The minimum RMSD_holo_ value across the holo structures (minRMSD_holo,X_) was then used as a measure of how closely the frame resembles a holo state. Only the binding site residues (Intersection of Holo-PPS-MD1-3 and Intersection of Holo-PR-MD1-3 residue sets in Table S1) were used both for the alignment and for the RMSD calculation.

For the PR-to-PPS SMD trajectory, a similarity measure was introduced to determine if a given frame was more similar to the PR or to the PPS holo states. The minRMSD_holo,X_ profiles calculated along the trajectory were first rescaled between 0 and 1 and their complement to 1 was calculated, yielding the similarity metric Similarity_holo,*X*_:

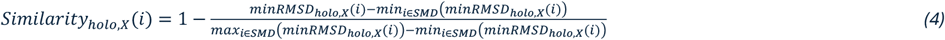

where *i* denotes an SMD frame and *X*=PR or PPS.

A given SMD frame was then defined as PR- or PPS-like by comparing its similarity values, with the frame assigned to the state *X* for which Similarity_holo,*X*_ was larger.

Next, we docked OM to the selected frames using AutoDock Vina^34,35^. OM was prepared with 8 rotatable bonds in total (including the 3 bonds recognised as amide type), and the frames from the sample100 and top100 sets were converted into pdbqt files using AutoDockFR^36^. A 20 × 20 × 20 Å^3^ docking grid with 1-Å spacing was placed in either the PPS or PR pocket centre, as defined before. The exhaustiveness parameter was set to 32, and a maximum number of 9 binding poses was generated. Docking calculations were run with five different random seeds. The top-ranked ligand pose from each run was selected for further analyses. The binding affinity was estimated by averaging the predicted binding affinity of the top-ranking poses from each run. Ligand RMSD values from a reference holo conformation (holo PPS energy-minimized structure for PPS apo-OM-docked complexes and holo PR MD1 energy-minimized structure for PR apo-OM-docked complexes) were calculated for each selected pose using spyRMSD^37^ after alignment of the binding pocket to the reference structure.

## Results

### Training and optimising the deep-learning model

To investigate the optimal training setting of our deep-learning (DL) model, we used part of the holo trajectories where myosin is in the PPS conformational state (holo-PPS-MD1-3) as our initial training dataset pool (n = 9,198) and the remaining holo-PPS-MD4 trajectory as the test dataset (n = 3,066). From the training dataset pool, we randomly sampled 600 frames for training and validation. A preliminary screening of different values for K-fold cross-validation (K = 3, 5, 6, 10) showed no significant impact of K on the results (analysis of variance, ANOVA, alpha = 0.05; Table S2). Based on this, we selected K = 3 as the default setting for subsequent model training and evaluation. Additionally, using a masked GDF loss function proved more effective than the standard GDF loss function (Figure S1).

Next, we examined the relationship between model accuracy and dataset size. We trained our models with varying dataset sampled sizes (size = 100, 300, 600, 1200, 2400, 4800). While models trained with larger datasets exhibited improved performance (Figure S2), when the dataset size exceeded 2,400 frames, the performance gains began to plateau, likely due to saturation of the model’s learning capacity (Table S3). On the other hand, smaller datasets (size <= 1,200) introduced sampling imbalances, which caused higher variability in cross-validation results. This variability decreased progressively as the dataset size increased, highlighting the importance of sufficient data for stable model training.

To ensure accurate predictions on test frames with arbitrary orientations (not necessarily aligned with the training frames), we applied 3D random rotations as data augmentation during training. This augmentation caused a small decrease in performance on non-rotated test frames compared with models trained without augmentation (Table S4), but it substantially improved performance on randomly rotated test frames (Figure S4). Among the evaluated rotation ranges, −180° to 180° provided the most consistent performance (Table S4). To reduce the performance drop associated with augmentation, we increased the training set to 4800 frames.

The optimal DL model (PPS-trained-4800 model) was trained with 3-fold cross-validation, a masked GDF loss function, a sampled holo-PPS dataset size of 4,800 frames and 3D random rotations with a rotation range from −180° to 180°. This model achieved mIoUs of 0.799 ± 0.002 and 0.771 ± 0.001 on the training and validation sets. The model with the highest validation mIoU was then tested on an unseen dataset (holo-PPS-MD4), achieving a test mIoU of 0.756 ± 0.096 (Figure S5).

Building upon the PPS-trained-4800 model, we extended its training to include a second conformational state of myosin, PR. This state is still able to bind OM, but the binding site composition and shape are different from the PPS state. We adopted a transfer learning approach, where we retrained the model starting from the PPS-trained-4800 weights and using a mixed dataset comprising 1600 frames randomly sampled from holo PPS (holo-PPS-MD1-3) and PR (holo-PR-MD1-3) trajectories. After 200 epochs of training (Figure S7), the new DL model (mix-1600-TL model) achieved a performance comparable to the previous model, with training and validation mIoUs of 0.793 ± 0.002 and 0.750 ± 0.003. In contrast, training with the same mixed dataset without transfer learning resulted in training and validation mIoUs of 0.726 ± 0.006 and 0.702 ± 0.002, respectively. These results underscore the effectiveness of transfer learning in maintaining high model performance, even with a reduced dataset size.

Finally, the model with the highest validation mIoU (0.752) was tested on unseen trajectories, holo-PPS-MD4 and holo-PR-MD4 (Figure 3), achieving a strong performance across both datasets (mIoU >= 0.760 and accuracy > 94%). These findings highlight the robustness of the mix-1600-TL model in generalising across different binding states and unseen datasets.

**Figure 3.**
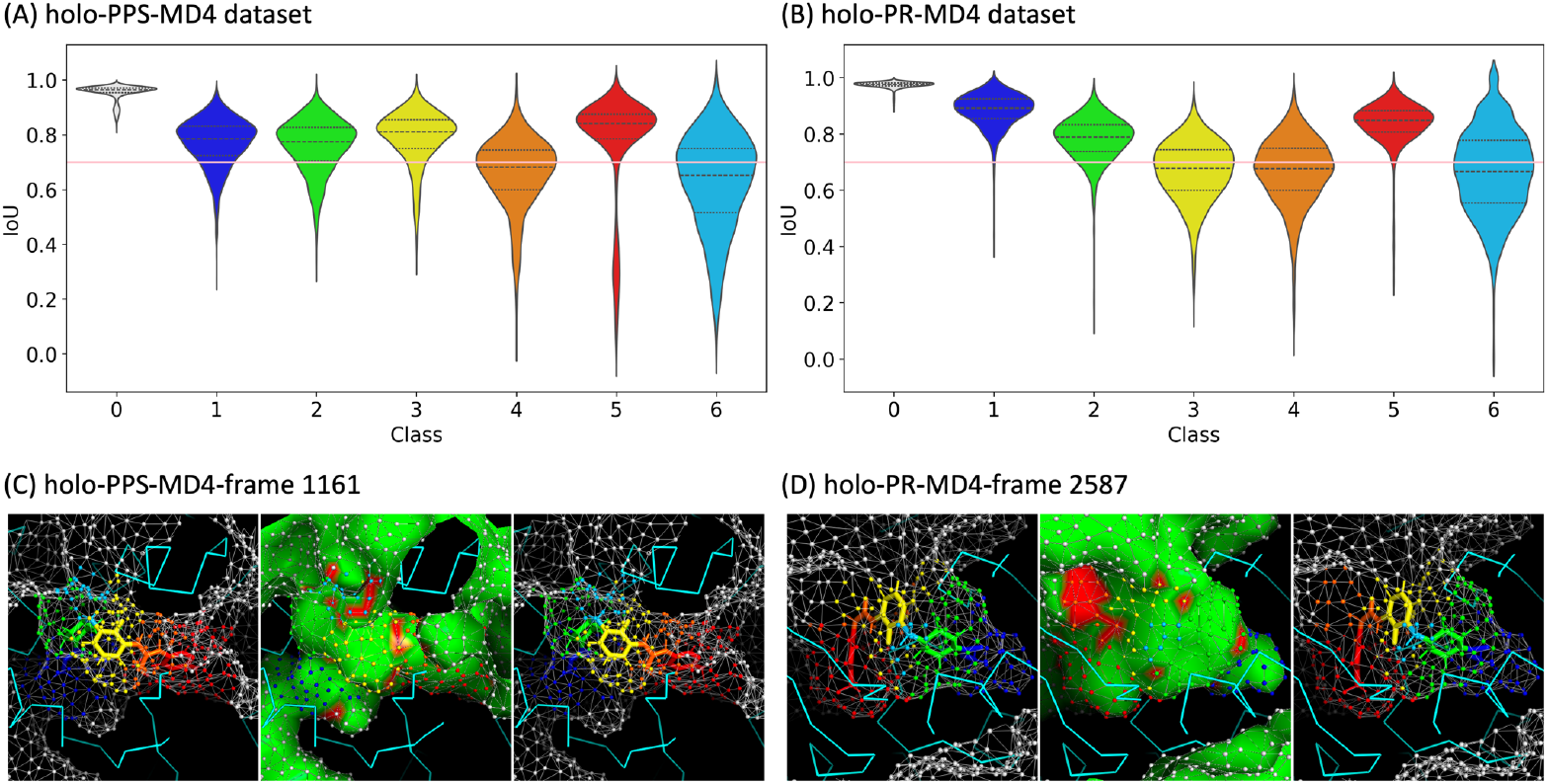
Performance of the optimal FragBEST-Myo (mix-1600-TL model) trained on a mixed PR and PPS dataset (sampled dataset size: 1600) with 3D random rotations (−180° to 180°). (A-B) Distributions of per-class IoU values obtained when the model was evaluated on the holo-PPS-MD4 (A) and holo-PR-MD4 (B) testing set (n=3066 for each set). Distributions are reported as violin plots (background class, c= 0), a horizontal pink line at 0.7 is included as a visual guide. (C-D) Analysis of model predictions for a sample frame from holo-PPS-MD4 (C, mIoU=0.760, accuracy=S4.8%, IoUs of classes from 0 to C: [0.971, 0.788, 0.780, 0.774, 0.630, 0.850, 0.531]) and from holo-PR-MD4 (D, mIoU=0.782, accuracy= S5.S%, IoUs of classes from 0 to C: [0.979, 0.8S8, 0.817, 0.67S, 0.323, 0.888, 0.889]). For each frame, the molecular surface (SES) in the ROI region is shown as mesh and coloured according to the ground truth (left panels) or predicted (middle and right) labels (same colours as panels A-B). Surface colouring in the middle panels indicates correct (green) and incorrect (red) predictions.

### Detection of holo-like forms in apo simulations

After assessing the DL model’s ability to segment binding sites in frames extracted from holo simulations, predictions from the mix-1600-TL model were converted into the “holo descriptors” D_1_-D_4_ to identify holo-like conformations in apo simulations. To enhance the conformational diversity of the samples considered for holo-like conformation detection, we analysed four different apo PPS trajectories (apo-PPS-MD1-4). The conformational distance from the holo trajectories used to train the DL model was evaluated by calculating the RMSD_holo_ values as described in the methods section.

To provide a benchmark, RMSD_holo_ and descriptor values were first calculated for the holo trajectory not used for model training (holo-PPS-MD4). RMSD_holo_ values ranged from 1 Å to 3 Å (Figure S8A-B), with the median value ≤ 2 Å for most of the holo-PPS-MD4 frames. The D_1_-D_4_ descriptors were calculated for each frame (Figure S8C-F, Table S5). D_1_ (number of distinct labels) remained constant at 7, indicating that all OM fragment types were assigned by the model to at least one vertex in all frames. D_2_ (fraction of vertices with any non-background label) ranged from 19.5% to 32.1%, with an average around 28%, while D_3_ (average fraction of vertices with a specific non-background label) ranged from 3.2% to 5.4%. A breakdown of D_3_ into individual fractions (Figure S8G) showed some variability across the different classes consistent with fragment size, with fragment 5 being the largest and fragment 6 the smallest. Interestingly, fragment 5 showed the strongest correlation with RMSD_holo_. D_4_ (averaged capped HoloSpace score) was found to be 1 for most of the frames, indicating sufficient pocket space to accommodate all fragments (Figure S8F). A clear correlation with RMSD_holo_ was observed for this descriptor, even if it does not use any information on the holo binding site volume.

The RMSD_holo_ and descriptor values were next calculated for the apo-PPS-MD1-4 trajectories and compared to the holo-PPS-MD4 benchmarks. RMSD_holo_ values ≤ 2 Å, indicating the presence of holo-like conformations, were observed in the apo-PPS-MD3 trajectory (Figure S9C) and more frequently in the apo-PPS-MD4 one (Figure S9D). Interestingly, apo frames were observed where the four descriptors (Figure 4A-D) fell within the expected range for holo-like forms (dark pink shading in the ‘holo zone’ panel). These frames were located more frequently in the low RMSD_holo_ parts of the trajectories. Accordingly, all descriptors exhibited anti-correlation with minRMSD_holo_ values (Figure S10), indicating that they contain useful information for identifying potential holo-like frames in the apo simulations.

**Figure 4.**
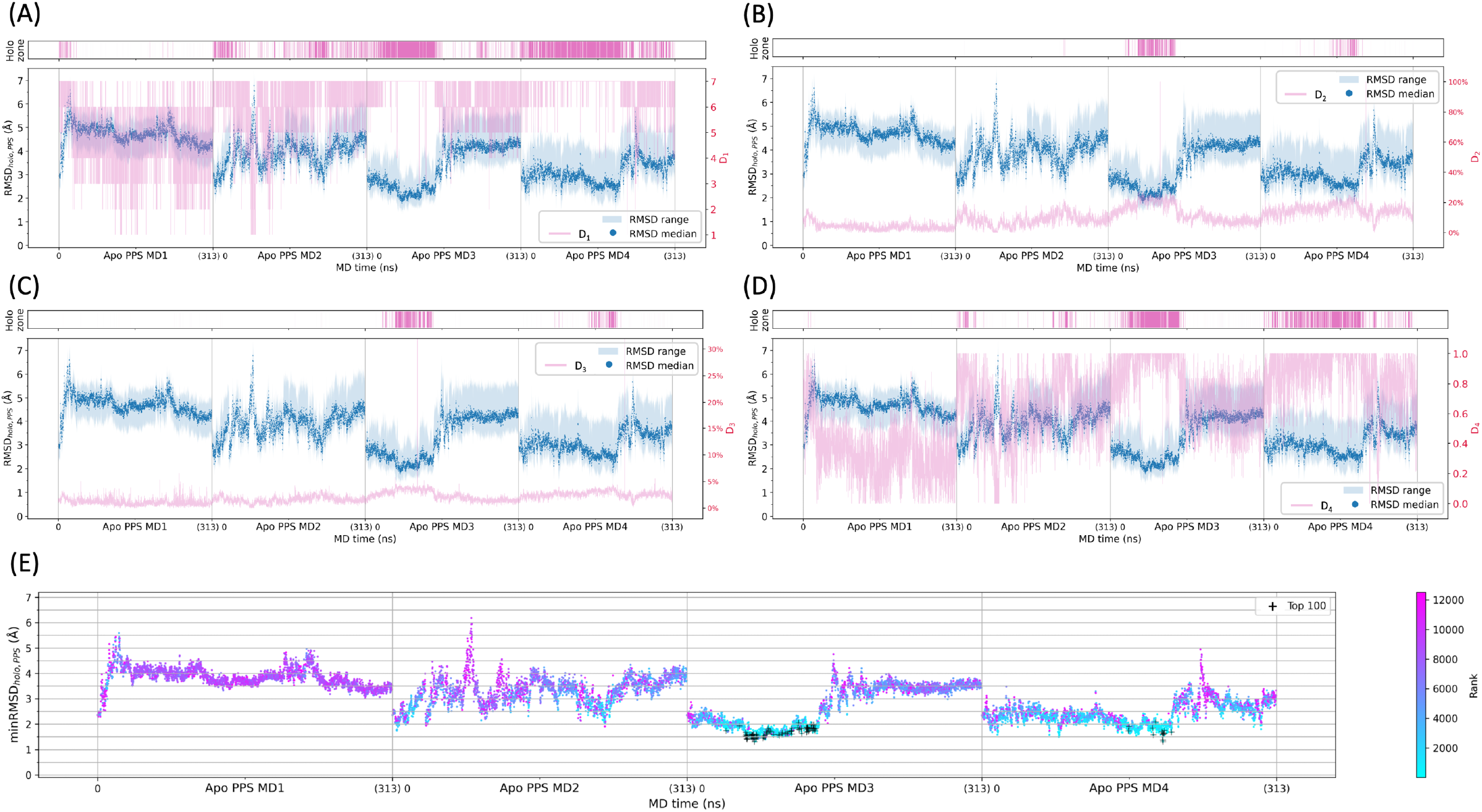
(A-D) Relationship between RMSD_holo,PPS_ values and deep-learning-derived descriptors D_1_-D_4_ (pink) for apo-PPS-MD1-4 simulations. The time evolution of RMSD_holo,PPS_ median values is shown in dark blue, with the min-max range indicated as light blue shading. Dark pink shading in the “holo zone” indicates frames with descriptor values within the holo-PPS-MD4 ranges (min-max values in Table S5). (E) Time evolution of minRMSD_holo,PPS_ values, where each point is coloured from cyan to magenta according to the R_holo_ rank of the frame. The top 100-ranked frames are indicated with a black cross.

All apo frames were then assigned a rank R_holo_ based on the D_1_-D_4_ descriptors (see Methods section). Higher-ranked frames (cyan in Figure 4E) were located again in the regions with lower minRMSD_holo_ values. Moreover, a positive association was found between ranking and minRMSD_holo_ (Figure S16A). Comparing apo frames with different R_holo_ ranks clearly showed surfaces segmented into multiple fragment-binding regions for the top-ranked frames (left panel in Figure 5A), while the number and extension of regions with non-background labels decreased with the rank (middle and right panel).

**Figure 5.**
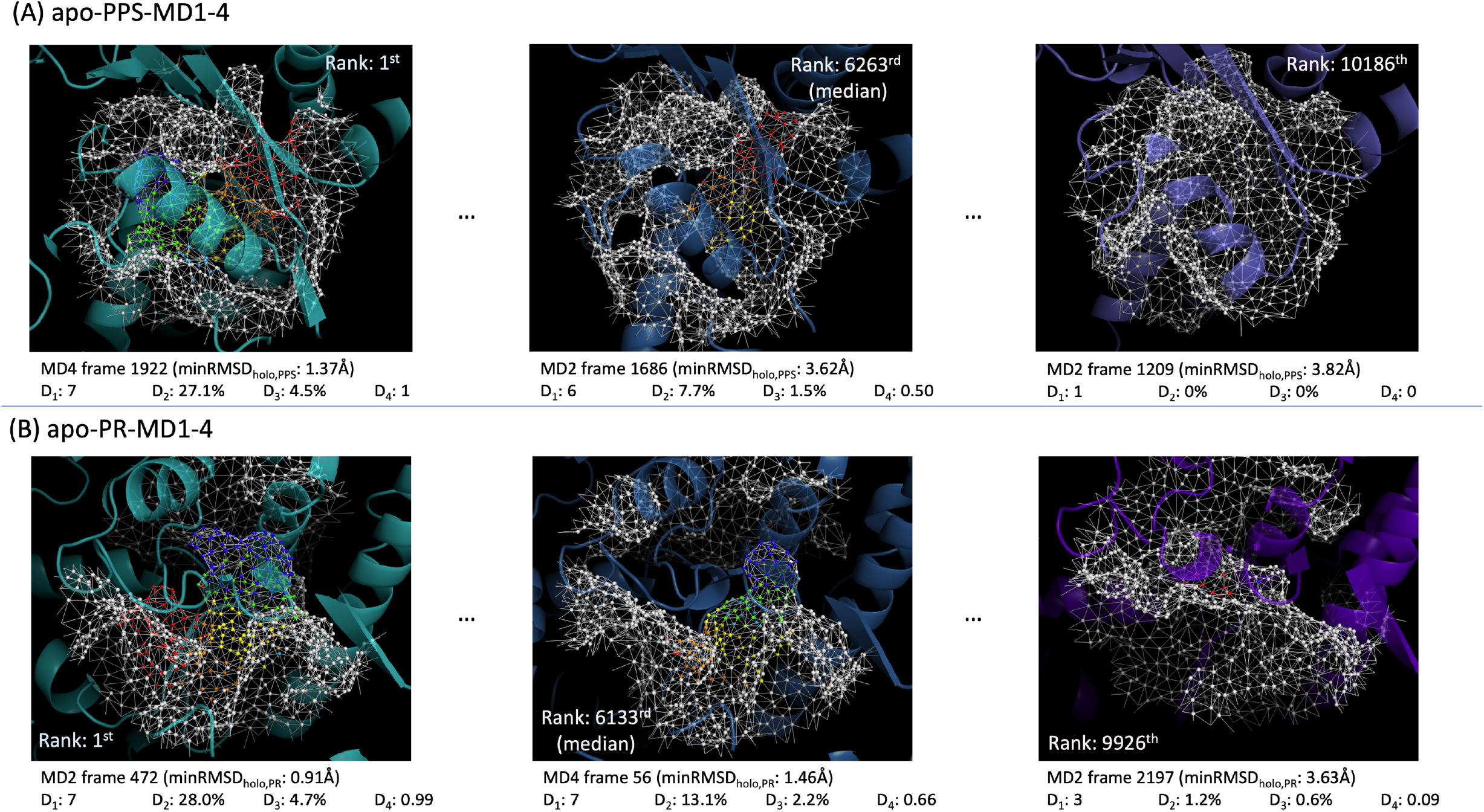
Frames selected from the top, middle and lower R_holo_ ranks of apo PPS (A) and PR (B) simulations. A cartoon representation is used for the protein, with the molecular surface in the ROI region shown as mesh. Vertices are coloured according to the label predicted by FragBest-Myo (see Figure 1B for the colour code, vertices labelled as background are shown in white).

Analyses of D_1_-D_4_ descriptors and R_holo_ ranks of PR simulations (Figures S11-S14, S16B) led to similar conclusions. Benchmark D_1_-D_4_ ranges observed in the PR holo simulations were comparable to the PPS ones (Table S5). The top-ranked apo frames were again found in low (≤ 2 Å) minRMSD_holo_ regions (Figure S13E) and featured more complex segmentation patterns, with a greater diversity of predicted labels, compared to frames with lower ranks (Figure 5B).

### Performance of FragBest-Myo-selected apo frames in ensemble docking

To test the use of R_holo_ ranking in ensemble docking, OM was docked to the top 100 ranking apo frames (top100) for each state (PPS and PR). Docking to 100 randomly selected frames (sample100) was also performed as a control. The resulting binding poses were then compared to reference poses from the holo simulations.

As expected from the analysis in the previous section, the top100 frames (orange in Figure 6A and S17A) showed significantly higher similarity to holo forms at the binding pocket compared to the sample100 (blue) ones. The binding poses obtained from docking OM to apo structures lack an exact reference for comparison, as native binding poses are observed in holo structures. Therefore, for each apo-docked structure, the closest reference was identified as the holo structure with the most similar binding site (minRMSD_holo_). The top100 docked frames showed consistently lower ligand RMSD values to the reference holo structure than the sample100 set, indicating that their closer structural similarity to holo binding sites also translated into better docking results. Specifically, 10% of top100 PPS frames achieved a ligand RMSD ≤ 3.5 Å compared to 0% of randomly selected conformations. Similarly, 19% of top100 PR frames had a ligand RMSD ≤ 3.5 Å compared to 4% in the random set (Figure 6C, S17C).

**Figure 6.**
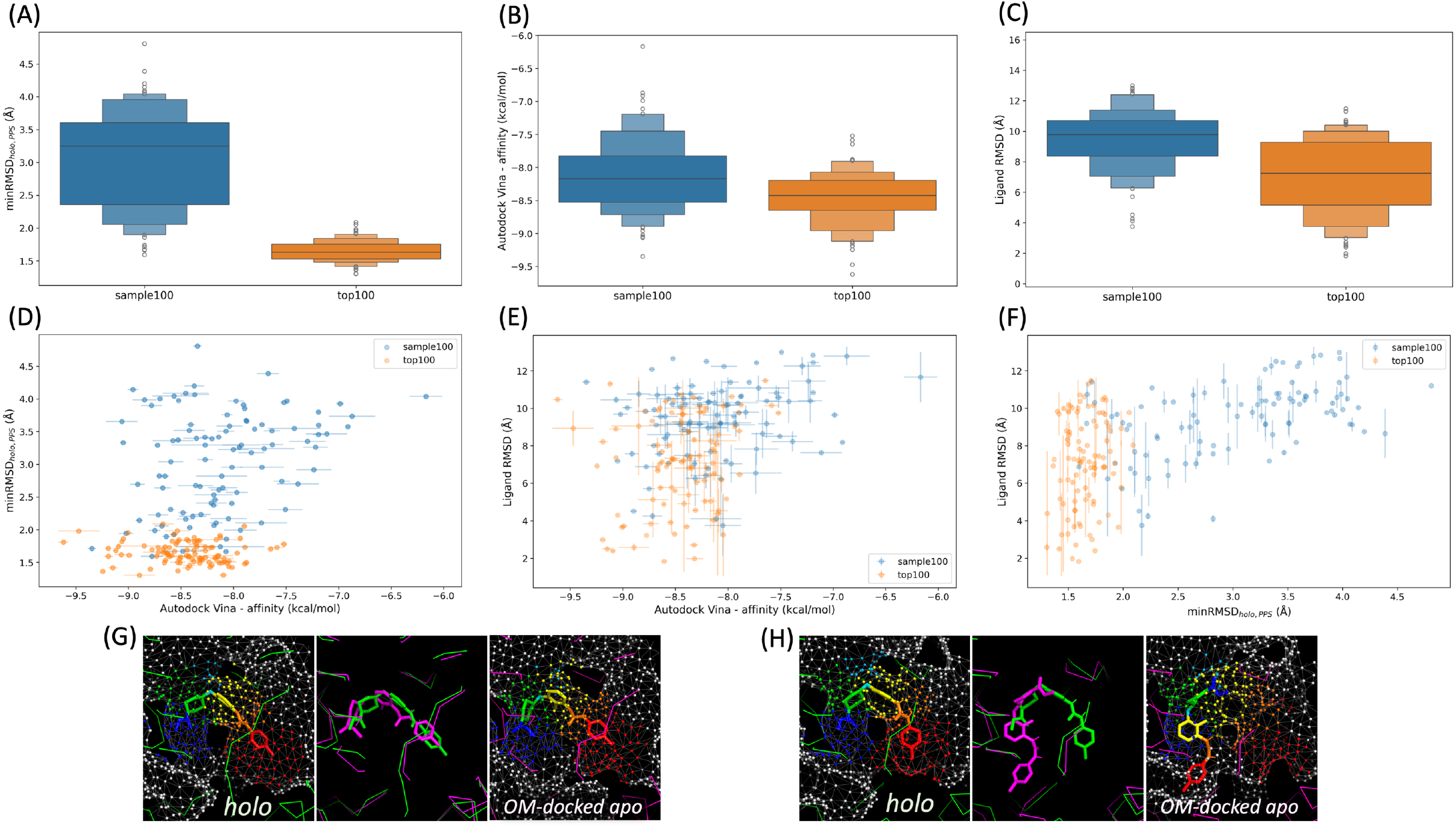
OM docking to selected frames from apo-PPS-MD1-4 trajectories. (A) Enhanced box plots of minRMSD_holo,PPS_ values for sample100 (blue) and top100 (orange) frames. (B) Enhanced box plots of Vina binding affinities of OM to sample100 and top100 frames. For each frame, the binding affinity is obtained by averaging the values calculated for the best docking poses from 5 runs. (C) Enhanced box plots of ligand RMSD values relative to the reference holo structure for OM docked to sample100 and top100 frames. RMSD values are calculated using heavy atoms only. (D-F) Scatter plots showing pairwise relationships between minRMSD_holo,PPS_, Vina binding affinity and ligand RMSD values to the reference holo structure. Error bars show standard deviations across 5 docking runs (using the best docking pose for each run). (G and H) Comparisons between selected OM-docked apo frames and their closest holo structure. The 57^th^-ranked frame (apo-PPS-MD3, frame 634) and its reference holo frame (holo-PPS-MD1, frame 2579) are shown in (G), while the 1_st_-ranked apo frame (apo-PPS-MD4, frame 1922) and its reference holo frame (holo-PPS-MD1, frame 2540) are shown in (H). Left panels: OM in the reference holo frame (sticks) coloured by fragments (see Figure 1B for the colour code), holo protein (green ribbon), protein surface mesh (white lines) and vertices with ground-truth labels (white indicates the background). Middle panels: superimposition of OM-docked apo (magenta) and reference holo (green) frames. Right panels: OM docked to the apo frame (sticks) coloured by fragments, apo protein (magenta ribbon), protein surface mesh, and vertices with predicted labels.

Using the top100 structures also resulted in stronger predicted binding affinities compared to the sample100 set (Figure 6B, S17B). Indeed, a correlation was observed between affinities and structural similarity to the holo references, both at the binding site (minRMSD_holo_) and in ligand poses (ligand RMSDs), with top100 frames (orange) generally clustering in the lower-left corner of the plots (Figure 6D-F, S17D-F).

A closer inspection of the top100 docked poses suggested that, in some cases, the mismatch between docked and reference poses was due more to docking itself than an incorrect frame selection. For illustration, two frames with high similarity to a holo frame (minRMSD_holo_ ~ 1.4 Å) and good agreement between the predicted (right panels in Figure 6G-H) and ground-truth (left panels) binding-site segmentation led to different docking outcomes. In one case, the OM docked pose was well aligned to the holo pose (Figure 6G, ligand RMSD: 2.41 Å, Vina affinity: −9.11 kcal/mol), while in the other OM was placed in a different region of the binding site (Figure 6H, ligand RMSD: 7.30 Å, Vina affinity: −8.54 kcal/mol). Importantly, the wrong docking pose was inconsistent with the predicted segmentation (right panel in Figure 6H), suggesting that deviations from FragBEST-Myo predictions can be used to detect non-native poses.

### Application to MD simulations sampling multiple states

After testing our method on MD trajectories that sample individual conformational states of myosin (PR or PPS), we applied it to a steered-MD (SMD) simulation of the myosin recovery stroke (PR to PPS transition), where both states are observed at different times. Due to the different composition and shape of the OM-binding pockets in the two states, we used two different ROI definitions based on the two different sets of binding site residues (Apo PR-to-PPS sets in Table S1). For each ROI definition, model predictions were used to calculate the descriptors and rank all frames in the SMD trajectory. In both cases, the top-ranked frames were found in low RMSD_holo_ regions (Figure 7). Consistent with the results from single-state simulations presented in the previous sections, the R_holo_ scores correlated with the similarity to the reference holo states (as measured by the Similarity_holo_ index), regardless of the ROI definition used for the predictions (Figure S18A-B).

**Figure 7.**
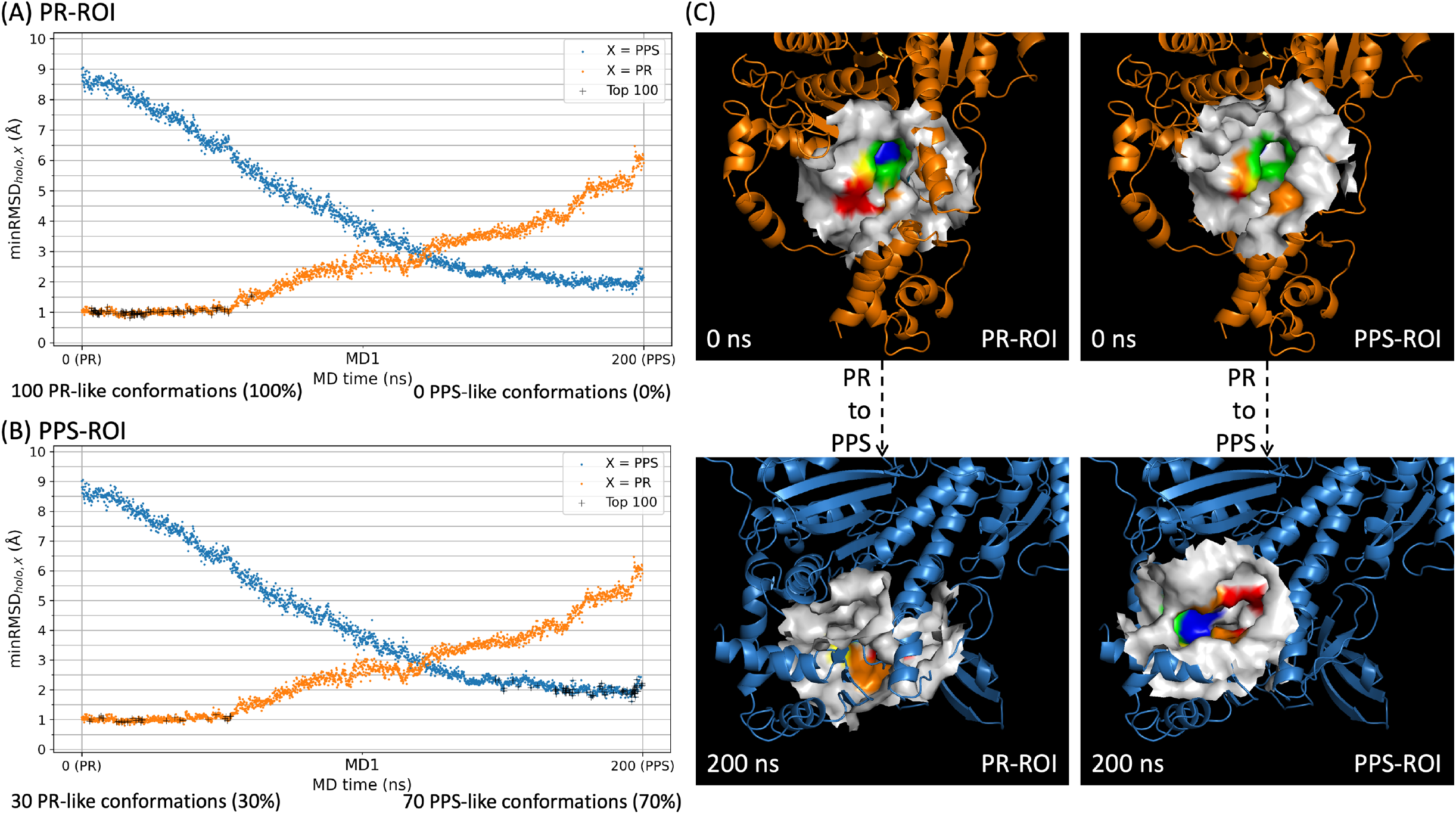
FragBEST-Myo predictions along the PR-to-PPS transition. (A-B) Time evolution of minRMSD_holo,X_ values during the apo PR-to-PPS SMD trajectory, with X=PR (orange) or PPS (blue). Black crosses indicate the top 100 frames ranked by FragBEST-Myo R_holo_ using the PR (A) or PPS (B) ROI definition. (C) Surface representation of the solvent-excluded surface in the PR (left panels) and PPS (right panel) ROIs used for the initial (t=0 ns) and final (t=200 ns) frames of the apo PR-to-PPS trajectory. The surface is coloured according to the predicted labels (see Figure 1B for the colour code, regions labelled as background are shown in white).

When using the PR ROI, all top100 frames had a minRMSD_holo,PR_ <= 2 Å (black crosses in Figure 7A). Interestingly, the PPS ROI definition led to selecting not only frames with low minRMSD_holo,PPS_ values (70% of top100 frames), but also frames close to the PR state (black crosses superimposed on the orange dots in Figure 7B). Inspection of the D_1_-D_4_ descriptors for the first and last frame of the trajectory shows that when using the PPS ROI, descriptor values fall within the reference holo ranges for both frames (Table 2). This is because the residue sets based on the PR and PPS pocket definitions have similar geometric centres in PR-like frames, meaning that PR-like pocket features can be captured using both ROI definitions (Figure 7C, upper panels). By contrast, the geometric centres of the two sets are far apart in PPS-like frames, so PPS-like pockets can be described correctly only using the PPS ROI (Figure 7C, lower panels). At the same time, using the PPS ROI triggered several warnings in the pre-ranking filtering stage, which resulted in a significant number of frames (~76.4%) being filtered out before ranking. This is in part related to the larger solvent-accessible space encompassed by the PPS-pocket ROI for a portion of the frames, which contributes to an overestimation of the HoloSpace used for the calculation of D_4_ (Figure S18C).

**Table 2.**
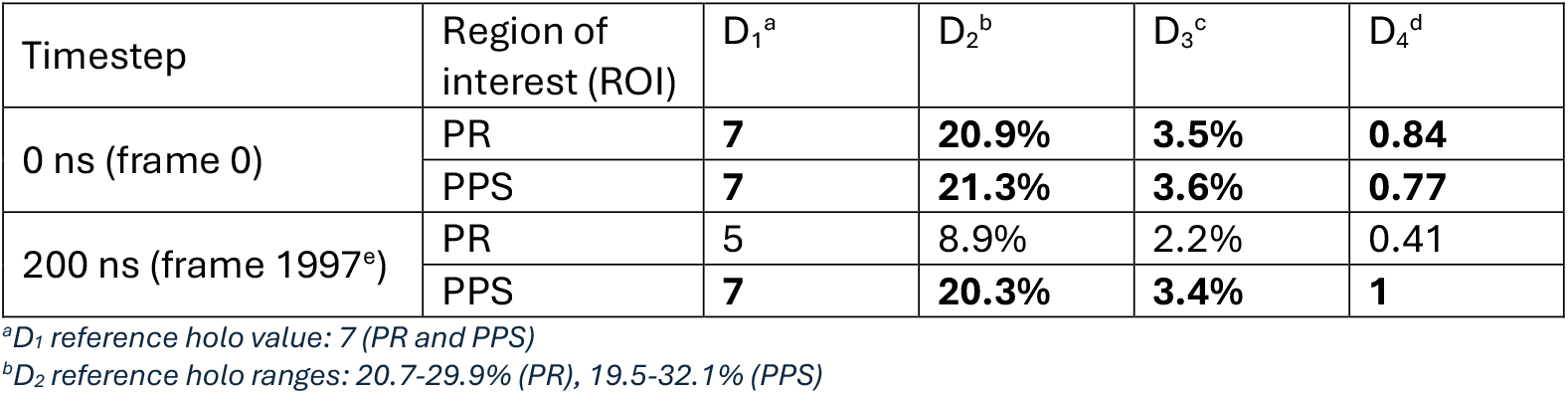
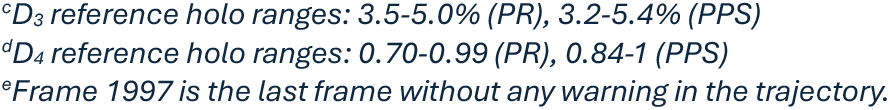
D_1_-D_4_ descriptors for the first and last frame of the PR-to-PPS SMD trajectory calculated using the PR and PPS ROI definition. Values within reference holo ranges are highlighted in bold.

| Timestep | Region of interest (ROI) | $D_1^a$ | $D_2^b$ | $D_3^c$ | $D_4^d$ |
| --- | --- | --- | --- | --- | --- |
| 0 ns (frame 0) | PR | **7** | **20.9%** | **3.5%** | **0.84** |
|  | PPS | **7** | **21.3%** | **3.6%** | **0.77** |
| 200 ns (frame 1997<sup>e</sup>) | PR | 5 | 8.9% | 2.2% | 0.41 |
|  | PPS | **7** | **20.3%** | **3.4%** | **1** |

<sup>a</sup> $D_1$ reference holo value: 7 (PR and PPS)
<sup>b</sup> $D_2$ reference holo ranges: 20.7-29.9% (PR), 19.5-32.1% (PPS)
<sup>c</sup>*D<sub>3</sub> reference holo ranges: 3.5-5.0% (PR), 3.2-5.4% (PPS)*
<sup>d</sup>*D<sub>4</sub> reference holo ranges: 0.70-0.99 (PR), 0.84-1 (PPS)*
<sup>e</sup>*Frame 1997 is the last frame without any warning in the trajectory.*

Next, we assessed the performance in ensemble docking of FragBEST-Myo-selected frames for both ROI definitions (top100_PR-ROI and top100_PPS-ROI) and compared it with our sample100 control (100 randomly sampled conformations from the apo PR-to-PPS trajectory). Each frame from all three sets was labelled as PR-like (Similarity_holo_,_PR_ > Similarity_holo_,_PPS_) or PPS-like (Similarity_holo_,_PPS_ > Similarity_holo_,_PR_), which resulted in 100% PR-like frames for top100_PR-ROI, 30% PR-like and 70% PPS-like frames for top100_PPS-ROI, and an almost equal representation of the two states in sample100 (46% PR-like and 54% PPS-like). Two docking grids (PR-grid and PPS-grid) were defined based on the corresponding PR and PPS ROI centres. Docking of FragBEST-Myo-selected frames was performed with the grid consistent with their ROI definition (PR-grid for top100_PR-ROI and PPS-grid for top100_PPS-ROI), while the sample100 control frames were docked using both grids. This setup reflects a scenario where binding pocket prediction tools have been used on apo structures representative of the PR and PPS states to identify potential binding sites, and the user wishes to test them both on all the sampled frames.

When compared to their controls (light shades in Figure 8A), FragBEST-Myo-selected frames (darker shades) showed much higher similarity to their corresponding holo forms. Consistently, OM docking to the top100 frames yielded better agreement with holo binding poses than sample100 (Figure 8C), with 21% (PR-like) and 7% (PPS-like) of top100 poses within 5 Å of the closest holo frame, versus 13% and 6% for sample100, respectively (Figure S19). Moreover, while distributions of top100 and sample100 predicted binding affinities were similar for PR-like frames (blue shades in Figure 8B), the PPS-like top100 distribution was shifted towards more negative values compared to its controls (orange shades). It is also worth noting that, based on both computational and experimental results, OM is expected to have a stronger binding affinity to the PPS state compared to the PR one^38^.

**Figure 8.**
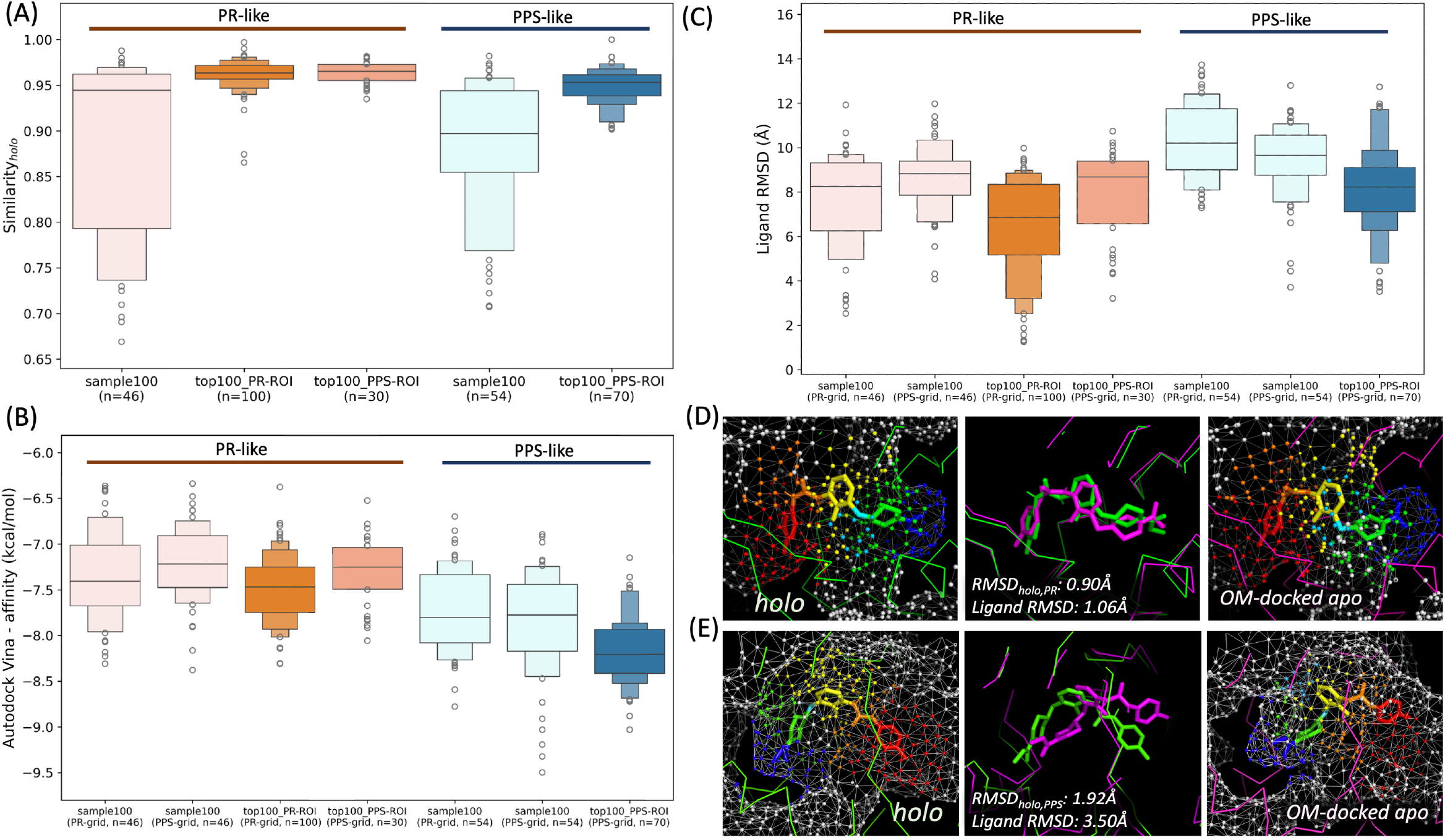
OM docking to selected frames from the apo PR-to-PPS SMD trajectory. (A) Enhanced box plots of Similarity_holo_ values comparing groups of PR-like (orange shades) and PPS-like (blue shades) frames. For each group, control results from the randomly selected set (sample100) are shown in the lightest shade. Results from the FragBEST-Myo selected frames (top100) are shown in darker shades, split by ROI location. The darkest shade corresponds to frames in states consistent with the ROI-location, with PR-like frames from top100_PR-ROI in darkest orange and PPS-like frames from top100_PPS-ROI in darkest blue. Intermediate orange denotes PR-like frames from the top100_PPS-ROI set. (B) Enhanced box plots of Vina binding affinities of OM to sample100 and top100 frames. For each frame, the binding affinity is obtained by averaging the values calculated for the best docking poses from 5 runs. The same colour code is used as for (A). The location of the docking grid used for the calculations is indicated in the x-axis label (PR-grid or PPS-grid). (C) Enhanced box plots of ligand RMSD values relative to the reference holo structure for OM docked to sample100 and top100 frames. RMSD values are calculated using heavy atoms only. The same colour code and labelling is used as for (B). (D and E) Comparisons between selected OM-docked apo frames and their closest holo structure. The 2^th^-ranked frame in top100_PR-ROI (frame 221, PR-like) and its reference holo frame are shown in (D), while the 17_th_-ranked frame in top100_PPS-ROI (frame 1739, PPS-like) and its reference holo frame are shown in (E). Left panels: OM in the reference holo frame (sticks) coloured by fragments (see Figure 1B for the colour code), holo protein (green ribbon), protein surface mesh (white lines) and vertices with ground-truth labels (white indicates the background). Middle panels: superimposition of OM-docked apo (magenta) and reference holo (green) frames. Right panels: OM docked to the apo frame (sticks) coloured by fragments, apo protein (magenta ribbon), protein surface mesh, and vertices with predicted labels.

When comparing the performance of PR-like and PPS-like FragBEST-Myo docked frames, the former showed better agreement with holo forms (Figures 8A and 8C). Indeed, it was possible to find PR-like top100 docking poses well superimposed on their closest holo frame (Figure 8D), while PPS-like poses, although correctly oriented, were less well aligned (Figure 8E). This is consistent with how the PR-to-PPS trajectory was generated^39^: starting from an experimental PR structure, the myosin lever arm was gradually rotated towards the PPS state without using any direct information on the PPS binding site. The starting point of the trajectory was thus inherently closer to the holo reference than the end point, and the relatively weaker performance of PPS-like frames should not be attributed to FragBEST-Myo limitations.

## Discussion

Here, we showed that a deep learning model (FragBEST-Myo) can be trained to recognise fragment-binding regions in apo frames from MD simulations of cardiac myosin, using holo simulations as training set. We also demonstrated that predictions from the model can be used to select apo frames more likely to produce holo-like docking poses of the cardiac activator OM than randomly selected frames.

Our initial analyses showed that segmentation performance depends on the number of training frames, with at least 600 frames required to achieve a test-set mIoU of ~0.7 and at least 2,400 frames needed to substantially reduce performance fluctuations across cross-validation folds. This indicates that it is essential to take into account protein dynamics when training the model.

Analysis of the D_1_-D_4_ descriptors calculated from the model predictions highlighted that similarity to holo-frames correlates with both the number and extent of segmented regions. An apo frame was found to be close to a holo one (minRMSD_holo_ ~ 2 Å or below) only when segmented regions were detected for all possible non-background classes (all OM fragments) and they were sufficiently large to allow fragment binding.

Our results also showed that a single model trained on both PPS and PR state simulations can identify holo-like frames in apo simulations where both states are sampled. The OM binding site has a completely different shape in these two states, with only partial overlap among the residues forming the binding interface. While this finding is based on a single protein system, it provides initial evidence that developing a unified FragBEST model able to accurately segment multiple, distinct binding sites is possible. In addition, our results suggest that the training of such model could be accelerated by transfer learning.

To assess the practical implications of these findings, we next evaluated whether the model predictions could be used in docking applications. A natural application of a general FragBEST tool would be the selection of frames for ensemble docking studies where ligands are docked against a target binding site, but no direct structural information on the holo state of the site is available. To explore whether predictions from the current FragBEST-Myo model contain sufficient information for this task, we compared the performance of apo frames selected by FragBEST-Myo against a random selection. FragBEST-Myo frames were more likely to produce OM binding poses closer to the native one and with stronger binding affinities.

While FragBEST-Myo or any generalised version to be developed in the future can facilitate the identification of holo-like frames, it should not be interpreted as a method for generating them^15^. The model operates by detecting holo-like conformations within a pre-existing ensemble of apo structures, independent of how those structures were obtained. Consequently, it can be integrated with any approach for sampling conformational space, including MD methods with or without enhanced sampling^40,41^, or emerging deep learning–based techniques^42,43^.

It is also important to note that binding site segmentation produced by our model has potential applications beyond selection of holo-like frames. Even when the correct docking pose was not recovered, the FragBEST-Myo segmentation, including shape, location and fragment labels of the identified regions, was still consistent with the native pose. This indicates that the model provides valuable information not only for selecting frames suitable for docking but also for evaluating the quality of the resulting binding poses. Moreover, the predicted binding-site segmentation of a general FragBEST model could, in principle, be used as a template to guide fragment-based drug design^44,45^.

Training and applying FragBEST models require specifying a region of interest. When the binding pocket location is unknown, binding pocket prediction tools^46,47^ can be used. In this study, the location of the OM binding site was already known from experimental structures. However, it is worth noting that the fpocket^48^ predictor (v2.0, default settings) identifies pockets encompassing the OM binding site as high-ranking sites when run on the initial apo structures from both the PPS and PR simulations, consistently placing them among the top three pockets within the region close to the converter.

The FragBEST-Myo model (including model parameters, the code for ranking and selecting the conformations, and data for an example of ranking and selecting conformations) is freely available from https://github.com/fornililab/FragBEST-Myo, together with Jupyter notebook tutorials illustrating how to run the model on MD trajectories or sets of experimental structures, and how to use the predictions to rank frames for holo-like structure selection. As the model was developed as proof-of-concept and trained specifically on cardiac myosin bound to OM, its generalisation capability would be expected to be limited. However, when applied to experimental structures of other myosin isoforms (e.g., skeletal) or of cardiac myosin bound to different ligands (e.g., mavacamten), the predictions are consistent with expectations: ranking structures by R_holo_ places OM-bound conformations highest, followed by mavacamten-bound, with apo structures ranked last. This indicates that the model already captures transferable features beyond its original training set.

In summary, FragBEST-Myo shows that deep learning can be used to obtain fragment-specific segmentations of protein binding sites that incorporates information on intrinsic structural variability due to protein dynamics. These segmentations can be applied to different tasks, such as the detection of holo-like conformations as shown in this work, assessment of docking poses, and guidance of fragment-based design. The FragBEST framework therefore provides a basis for developing future models applicable to a broader range of proteins and ligands.

## Acknowledgment

This work has been supported by the BBSRC/QMUL AI for Drug Discovery Collaborative Training Partnership and BBSRC/QMUL Impact Acceleration Account Programme [grant no. BB/X511067/1], and used time on HPC granted via the UK High-End Computing Consortium for Biomolecular Simulation, HECBioSim (http://hecbiosim.ac.uk), supported by the EPSRC [grant no. EP/X035603/1]. Special thanks to Christopher Duffy, Harikrishna Jayanthan (former employee in Evotec), Alina Miron, Alessandro Pandini, Ping-Chung Yu, and members of the Fornili’s group for their valuable feedback provided at different stages of the project.

## Data and code availability

The code repository will be available at https://github.com/fornililab/FragBEST-Myo once this article is published. The example data and model parameters used in the provided tutorials in the code repository can be downloaded from Zenodo at https://zenodo.org/records/18630776.

## Author contributions

Conceptualisation: Arianna Fornili, Yu-Yuan Yang.

Project administration: Arianna Fornili, Yu-Yuan Yang.

Data curation: Yu-Yuan Yang.

Software development: Yu-Yuan Yang.

Visualisation: Yu-Yuan Yang.

Writing – original draft: Yu-Yuan Yang, Arianna Fornili.

Writing – review & editing: Yu-Yuan Yang, Arianna Fornili, Richard W. Pickersgill.

Funding acquisition: Arianna Fornili, Richard W. Pickersgill.

Supervision: Arianna Fornili, Richard W. Pickersgill.

