## Supplementary Information for "Learning fragment-based segmentation of binding sites from molecular dynamics: a proof-of-concept on cardiac myosin"

### Supplementary Methods

#### Model performance optimisation

##### *Choice of loss function*

A standard and a masked version of the combined generalised dice and focal loss (GDF) function were tested (main Methods). Learning curves showed a steady improvement in performance during the initial training phase until a plateau was reached (Figure S1). Training with the GDF loss function (A) required more epochs to reach the plateau compared to the masked GDF loss (B).

##### *Dataset size and sampling*

We examined the relationship between the model accuracy and dataset size by randomly sampling training and validation datasets from the training dataset pool (holo-PPS-MD1-3 set of replicas in Table S1). Different sample sizes (100, 300, 600, 1200, 2400, 4800) were considered. To evaluate the overall performance of models under different training settings, the best training and validation mIoUs were averaged across three folds of cross-validation. The model with the highest validation mIoU was then evaluated using the testing set, where mIoU was calculated for each frame and averaged across all frames. Table S3 summarises the results obtained with different dataset sizes, using fixed random seeds to split the dataset and initialise the model weights to ensure reproducibility. Experiments conducted with different random seeds yielded similar trends.

In general, larger sampled training datasets improved both training and validation mIoU (Figure S2). The reduced standard deviation in the best training and validation mIoU for sample sizes greater than 2400 suggests that variability due to random data sampling is minimal for these sample sizes. While test mIoU values did not improve significantly for sample sizes larger than 600, we still chose 2400 as baseline size to improve robustness of models and reduce their sensitivity to data splitting. During the subsequent training, we monitored the standard deviation of mIoU in the training and validation sets to mitigate bias from data sampling and increased the dataset size when needed (e.g., with data augmentation as described below).

The best FragBEST-Myo model trained with a dataset size of 2400 (sampled from the holo-PPS-MD1-3 set of replicas) achieved an overall test mIoU of 0.762 ± 0.102. The median per-class IoU exceeded 0.7 for all classes except 4 and 6 (Figure S3A). Visualisation of a representative test frame (with an mIoU value close to the test set average) revealed consistent and acceptable prediction error patterns (Figure S3B-D), where inaccuracies were most prominent at boundaries between regions with different predicted labels. Additionally, classes 4 and 6 correspond to smaller fragments, leading to smaller labelled regions with fewer vertices. This makes their IoUs more susceptible to boundary shifts.

These factors likely contributed to the higher error rate observed for these classes, highlighting the influence of fragment size and vertex count on model performance.

#### Training of the multi-state model

Preliminary tests showed that both the optimal PPS-trained-4800 model and a second model trained with the same protocol on the holo-PR-MD1-3 trajectories using PPS-trained-4800 weights (PR-trained-4800) have poor zero-shot performance when applied to the opposite state. The PPS-trained-4800 model achieved a very low mIoU when tested on four holo-PR trajectories (0.114 at best, Figure S6A), while the PR-trained-4800 model performed well on the holo PR test trajectory (mIoU: 0.750 ± 0.052) but dropped when tested on the holo-PPS-MD4 trajectory (mIoU of 0.133 ± 0.013, Figure S6B). This outcome was expected, as the binding pocket in the two states differed substantially in shape and residue composition due to significant conformational changes in the OM binding region. Inspecting the class-wise IoU distribution shows that high values were only obtained for the background class ($c$=0). Interestingly, some classes (3, 4 and 5) showed some transferability across states albeit limited.

Based on this results, to obtain an enhanced model that can handle both PPS and PR states, we re-trained the PPS-trained-4800 model via transfer learning (PPS-trained-4800 weights were used as starting point) on a mixed dataset of 1600 frames randomly sampled from both holo PPS (holo-PPS-MD1-3) and PR (holo-PR-MD1-3) trajectories (with masked GDF loss function and 3D-rotated data augmentation using a rotation range of -180° to 180°). After 200 epochs of training (Figure S7), the new DL model (mix-1600-TL) achieved performance comparable to the best single-state models (training mIoU: 0.793 ± 0.002; validation mIoU: 0.750 ± 0.003. Training on the same mixed dataset from scratch resulted in a lower performance (training mIoU: 0.726 ± 0.006; validation mIoU: 0.702 ± 0.002), highlighting the advantages of transfer learning when smaller datasets are used.

On independent test trajectories (Figure 3), mix-1600-TL performed well on both holo-PPS-MD4 and holo-PR-MD4, achieving mIoUs of 0.760 ± 0.088 and 0.782 ± 0.060 and accuracies of 94.1% ± 3.2% and 95.9% ± 1.4%, respectively.

#### Pre-ranking filters

After calculating the $D_{1-4}$ descriptors and before ranking frames, filters were applied to remove frames with potential anomalies in the number or label distribution of ROI vertices.

For the holo PPS (Figure S8H) and PR (Figure S11H) trajectories, vertex counts were generally stable, with typical ranges between 1000 and 1200. In contrast, isolated frames from the apo trajectories showed large drops: for example, two frames from the apo PPS trajectories contained 5 ROI vertices or less, which was traced to errors in MSMS mesh generation and the triangulation workflow. Because in these cases the vertices cover only a very small part of the binding site and would lead to unreliable descriptor values, a filter was applied so that all frames with 200 ROI vertices or less were removed before ranking. This issue was rare, affecting only 0.016% and 0.008% of apo PPS and PR frames, respectively.

A second potential source of anomalies was investigated by assessing the spatial distribution of vertices assigned to the same predicted class. In some cases, same-class vertices were found in regions that were far apart, which is less consistent with well-defined fragment-binding regions (expected to be relatively compact) and can potentially lead to unrealistically large HoloSpace volumes. To address this, we performed clustering analyses within each set of same-class vertices using DBSCAN[1,2]. The parameters were selected by grid-search: $\varepsilon$ (neighbourhood radius) was set to 5 Å, and minPts (minimum number of points in the $\varepsilon$-neighbourhood) was set to 1/3 of the number of vertices being clustered. Vertices recognised as outliers were then excluded from HoloSpace calculations (e.g., class 3 vertices in Figure S15). In some cases, DBSCAN identified multiple clusters of comparable size within a given set of same-class vertices, so outlier removal did not isolate a single dominant contiguous region and the resulting raw HoloSpace remained unrealistically large (e.g., class 6 vertices in Figure S15). A further filter was then applied to exclude frames with excessively large HoloSpace regions even after DBSCAN processing. Specifically, frames were removed with capped HoloSpace Volume ($cHSV_c$) for class $c$ larger than 10 times the RDKit volume of the corresponding class. This threshold was based on the $cHSV_c$ distributions observed for the holo trajectories (Figure S8I and S11I).

It is to be noted that, while the results presented in this work were obtained by applying these filters, the code distributed with this manuscript does not automatically remove frames. Instead, it raises warnings, allowing users to decide whether to remove the flagged frames or to further investigate them.

#### Supplementary tables

*Table S1. Pocket residues, dataset class, and number of frames for each MD replica or set of MD replicas (PPS: pre-power stroke state, PR: post-rigor state, SMD: steered MD).*

| MD replicas or set of MD replicas | Pocket residues (by resid) | Dataset class | # Frames |
|---|---|---|---|
| Holo-PPS-MD1 | 120, 146-148, 160, 163-164, 167-168, 492, 497, 500, 666-667, 710-713, 721-722, 762, 765, 767, 770-771, 774 | Training/Validation | 3066 |
| Holo-PPS-MD2 | 120, 146-147, 160, 163-164, 167-168, 492, 497, 500, 666, 710-713, 721-722, 762, 765, 770-771, 774 | Training/Validation | 3066 |
| Holo-PPS-MD3 | 120, 146-147, 160, 163-164, 167-168, 492, 497, 500, 666-667, 710-713, 721-722, 762, 765, 767, 770-771, 774 | Training/Validation | 3066 |
| Intersection of Holo-PPS-MD1-3 | 120, 146-147, 160, 163-164, 167-168, 492, 497, 500, 666, 710-713, 721-722, 762, 765, 770-771, 774 | - | - |
| Holo-PPS-MD4 | 120, 146-148, 160, 163-164, 167-168, 492, 497, 500, 666, 710-713, 721-722, 762, 765, 770-771, 774 | Testing (model performance) | 3066 |
| Holo-PR-MD1 | 84, 89-94, 96, 101, 118-121, 489, 493, 497, 500, 697-699, 701-702, 705, 709-714, 762 | Training/Validation | 3066 |
| Holo-PR-MD2 | 84, 89-94, 96, 101, 118-121, 493, 496-497, 500, 698, 701-702, 705, 709-713, 762, 770 | Training/Validation | 3066 |
| Holo-PR-MD3 | 84, 89-94, 96, 101, 118-121, 493, 497, 698, 701-702, 705, 709-713, 762, 770 | Training/Validation | 3066 |
| Intersection of Holo-PR-MD1-3 | 84, 89-94, 96, 101, 118-121, 493, 497, 698, 701-702, 705, 709-713, 762 | - | - |
| Holo-PR-MD4 | 84, 90-94, 96, 101, 118-121, 493, 497, 698, 701-702, 705, 709-713, 762, 770 | Testing (model performance) | 3066 |
| Apo-PPS (MD1-MD4) | 120, 146-147, 160, 163-164, 167-168, 170, 492, 497, 666-667, 710-713, 721-722, 765, 770, 771, 774 | Testing (holo-like frames detection) | 3131*4 |
| Apo-PR (MD1-MD4) | 84, 89-94, 96, 101, 118-121, 489, 493, 497, 500, 697-699, 701-702, 704-705, 710-713, 722, 762, 765, 770 | Testing (holo-like frames detection) | 3066*4 |
| Apo PR-to-PPS (SMD) | PPS pocket: 120, 146-147, 160, 163-164, 167-168, 170, 492, 497, 666-667, 710-713, 721-722, 765, 770-771, 774 | Testing (holo-like frames detection) | 2001 |
| | PR pocket: 84, 89-94, 96, 101, 118-121, 489, 493, 497, 500, 697-699, 701-702, 704-705, 710-713, 722, 762, 765, 770 | | |

*Table S2. Model performance for different cross-validation K values with a dataset size of 600. The dataset was sampled from the holo-PPS-MD1-3 set of replicas. No significant difference in the performance obtained using different K values was observed (using analysis of variance, ANOVA, alpha=0.05).*

| K | Train/validation set size | Composition (MD1/MD2/MD3)[a] | K-fold cross-validation | | Runtime (1x NVIDIA GeForce RTX2080) |
|---|---|---|---|---|---|
| | | | Best train mIoU[b] | Best validation mIoU[b] | |
| 3 | 400/200 | 233/166/211 | 0.900 ± 0.042 | 0.748 ± 0.031 | ~28hr |
| 5 | 480/120 | | 0.895 ± 0.076 | 0.750 ± 0.051 | ~50hr |
| 6 | 500/100 | | 0.810 ± 0.194 | 0.677 ± 0.151 | ~60hr |
| 10 | 540/60 | | 0.922 ± 0.109 | 0.770 ± 0.077 | ~105hr |

[a] *The dataset composition is reported as number of frames sampled from each replica.*

[b] *Train and validation mIoU values are reported as averages and standard deviations over the best per-fold values.*

*Table S3. Model performance with different dataset sizes. The datasets were sampled for the holo-PPS-MD1-3 set of replicas.*

| Sample size[a] | Train/validation set size | Composition (MD1/MD2/MD3)[b] | 3-fold cross-validation | | Test mIoU[d] |
|---|---|---|---|---|---|
| | | | Best train mIoU[c] | Best validation mIoU[c] | |
| 100 | ~67/~33 | 29/42/29 | 0.543 ± 0.089 | 0.446 ± 0.062 | 0.477 ± 0.057 |
| 300 | 200/100 | 90/112/98 | 0.711 ± 0.038 | 0.610 ± 0.036 | 0.608 ± 0.078 |
| 600 | 400/200 | 187/212/201 | 0.727 ± 0.137 | 0.635 ± 0.099 | 0.723 ± 0.096 |
| 1200 | 800/400 | 382/425/393 | 0.770 ± 0.142 | 0.650 ± 0.108 | 0.736 ± 0.098 |
| 2400 | 1600/800 | 804/810/786 | 0.968 ± 0.002 | 0.818 ± 0.001 | 0.762 ± 0.102 |
| 4800 | 3200/1600 | 1615/1633/1552 | 0.960 ± 0.002 | 0.824 ± 0.001 | 0.757 ± 0.107 |

[a]*Larger datasets included smaller ones (e.g., the 2400-size dataset contains all frames in the 1200-size dataset plus 1200 additional sampled frames). Frame counts were better balanced across the MD1, MD2 and MD3 replicas for larger dataset sizes.*
[b]*The dataset composition is reported as number of frames sampled from each replica.*
[c]*Train and validation mIoU values are reported as averages and standard deviations over the best per-fold values.*
[d]*The model with the highest validation mIoU across the folds was used for testing. The test mIoU values are reported as averages and standard deviations over all frames from the testing set (holo-PPS-MD4).*

*Table S4. Performance of the models trained on a 2400-frame sample of the holo-PPS-MD1-3 set of replicas with and without data augmentation (3D random rotation).*

| Rotation angle range | 3-fold cross-validation | | | Test mIoU[b] |
|---|---|---|---|---|
| | Best train mIoU[a] | Best validation mIoU[a] | Best model | |
| No rotation | 0.968 ± 0.002 | 0.818 ± 0.001 | Kfold 3, epoch 191, mIoU=0.818 | 0.762 ± 0.102 |
| -45° to 45° | 0.827 ± 0.003 | 0.793 ± 0.003 | Kfold 1, epoch 193, mIoU=0.796 | 0.762 ± 0.093 |
| -90° to 90° | 0.786 ± 0.001 | 0.763 ± 0.002 | Kfold 3, epoch 199, mIoU=0.764 | 0.752 ± 0.091 |
| -180° to 180° | 0.783 ± 0.001 | 0.758 ± 0.001 | Kfold 2, epoch 198, mIoU=0.759 | 0.747 ± 0.098 |

[a]*Train and validation mIoU values are reported as averages and standard deviations over the best per-fold values.* [b]*The model with the highest validation mIoU across the folds was used for testing. The test mIoU values are reported as averages and standard deviations over all frames from the testing set (holo-PPS-MD4).*

*Table S5. Summary of $D_{1-4}$ descriptor values observed during PPS and PR holo and apo trajectories. Mean values are reported with their standard deviation.*

| Trajectory set | $D_1$[a] | $D_2$ | Mean fraction of ROI vertices for each class | $D_3$ | $D_4$ | Mean *cHSS*[b] | Outliers[c] |
|---|---|---|---|---|---|---|---|
| Holo-PPS-MD4<br>(n = 3066) | 1: 0 (0%)<br>2: 0 (0%)<br>3: 0 (0%)<br>4: 0 (0%)<br>5: 0 (0%)<br>6: 0 (0%)<br>7: 3066 (100.0%) | Mean: 28.1 ± 1.5%<br>Min: 19.5%<br>Max: 32.1%<br>Median: 28.2% | 0: 71.9 ± 1.5%<br>1: 5.7 ± 0.6%<br>2: 4.6 ± 0.5%<br>3: 4.5 ± 0.5%<br>4: 3.5 ± 0.4%<br>5: 8.0 ± 1.0%<br>6: 1.8 ± 0.4% | Mean: 4.7 ± 0.2%<br>Min: 3.2%<br>Max: 5.4%<br>Median: 4.7% | Mean: 1.00 ± 0.02<br>Min: 0.84<br>Max: 1.00<br>Median: 1.00 | 1: 1.00 ± 0.00<br>2: 1.00 ± 0.01<br>3: 1.00 ± 0.01<br>4: 1.00 ± 0.00<br>5: 1.00 ± 0.00<br>6: 0.98 ± 0.10 | No |
| Apo-PPS-MD1-4<br>(n = 3131*4) | 1: 35 (0.3%)<br>2: 179 (1.4%)<br>3: 438 (3.5%)<br>4: 1003 (8.0%)<br>5: 1678 (13.4%)<br>6: 2844 (22.7%)<br>7: 6347 (50.7%) | Mean: 10.1 ± 6.2%<br>Min: 0.0%<br>Max: 100.0%<br>Max (excluding outliers): 27.6%<br>Median: 9.3% | 0: 89.9 ± 6.2%<br>1: 1.4 ± 1.6%<br>2: 1.6 ± 1.5%<br>3: 2.0 ± 1.4%<br>4: 2.2 ± 2.0%<br>5: 1.9 ± 1.8%<br>6: 0.9 ± 0.7% | Mean: 1.9 ± 1.3%<br>Min: 0.0%<br>Max: 100.0%<br>Max (excluding outliers): 6.6%<br>Median: 1.8% | Mean: 0.65 ± 0.27<br>Min: 0.00<br>Max: 1.00<br>Median: 0.68 | 1: 0.53 ± 0.47<br>2: 0.60 ± 0.45<br>3: 0.76 ± 0.38<br>4: 0.73 ± 0.41<br>5: 0.66 ± 0.45<br>6: 0.63 ± 0.43 | Yes<br>(MD3 frame 1068, MD4 frame 2164) |
| Holo-PR-MD4<br>(n = 3066) | 1: 0 (0%)<br>2: 0 (0%)<br>3: 0 (0%)<br>4: 0 (0%)<br>5: 0 (0%)<br>6: 0 (0%)<br>7: 3066 (100.0%) | Mean: 24.9 ± 1.3%<br>Min: 20.7%<br>Max: 29.9%<br>Median: 24.9% | 0: 75.1 ± 1.3%<br>1: 5.3 ± 0.8%<br>2: 4.8 ± 0.5%<br>3: 4.0 ± 0.6%<br>4: 3.0 ± 0.5%<br>5: 7.0 ± 0.6%<br>6: 0.9 ± 0.2% | Mean: 4.2 ± 0.2%<br>Min: 3.5%<br>Max: 5.0%<br>Median: 4.2% | Mean: 0.83 ± 0.02<br>Min: 0.70<br>Max: 0.99<br>Median: 0.83 | 1: 0.98 ± 0.07<br>2: 1.00 ± 0.00<br>3: 1.00 ± 0.00<br>4: 1.00 ± 0.01<br>5: 1.00 ± 0.00<br>6: 0.03 ± 0.08 | No |
| Apo-PR-MD1-4<br>(n = 3066*4) | 1: 0 (0%)<br>2: 1 (0.0%)<br>3: 41 (0.3%)<br>4: 810 (6.6%)<br>5: 1271 (10.4%)<br>6: 2241 (18.3%)<br>7: 7899 (64.4%) | Mean: 15.9 ± 5.6%<br>Min: 1.2%<br>Max: 29.1%<br>Median: 16.2% | 0: 84.1 ± 5.7%<br>1: 3.2 ± 2.7%<br>2: 2.9 ± 2.0%<br>3: 2.4 ± 1.3%<br>4: 2.2 ± 1.1%<br>5: 4.3 ± 2.0%<br>6: 0.8 ± 0.5% | Mean: 2.9 ± 0.8%<br>Min: 0.4%<br>Max: 7.7%<br>Median: 3.0% | Mean: 0.74 ± 0.17<br>Min: 0.00<br>Max: 1.00<br>Median: 0.81 | 1: 0.63 ± 0.45<br>2: 0.69 ± 0.43<br>3: 0.89 ± 0.30<br>4: 0.94 ± 0.20<br>5: 0.94 ± 0.20<br>6: 0.36 ± 0.37 | Yes<br>(MD1 frame 2380) |

[a]*Number and percentage of frames with a given $D_1$ value.*

[b]*Mean capped HoloSpace score with standard deviation for each non-background class.*

[C]*Presence of frames with less than 200 ROI vertices.*

#### Supplementary figures

(A) With GDF loss (B) With masked GDF loss

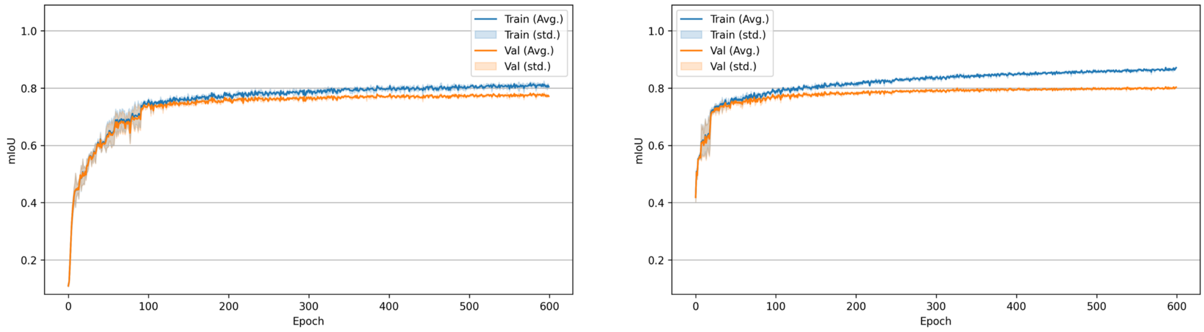


*Figure S1. Model training progress over 600 epochs when using standard (A) and masked (B) GDF loss. Models were trained on a 2400-frame sample of the holo-PPS-MD1-3 dataset with 3-fold cross-validation. The training (blue line) and validation (orange line) mIoU values averaged over the folds are reported for each epoch, together with their standard deviation (shaded area). The best training mIoU values are 0.819 ± 0.004 (A) and 0.873 ± 0.002 (B), while the best validation mIoU values are 0.785 ± 0.000 (A) and 0.805 ± 0.002 (B).*

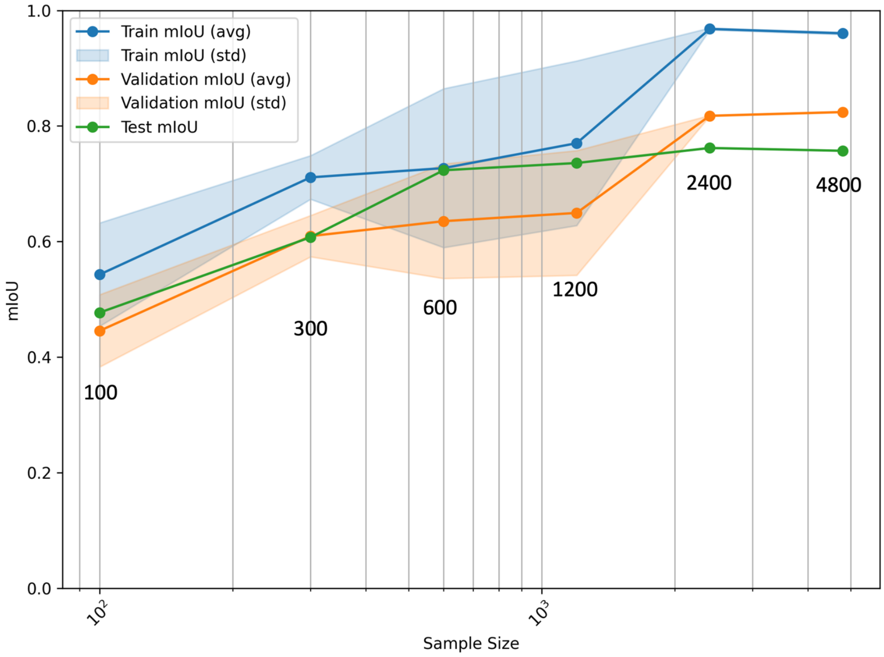


*Figure S2. Performance of models trained with different sampled dataset sizes and 3-fold cross validation. Datasets were sampled from the holo-PPS-MD1-3 set of replicas. The training (blue) and validation (orange) best mIoU values averaged over the folds are reported, together with their standard deviation (shaded area).*

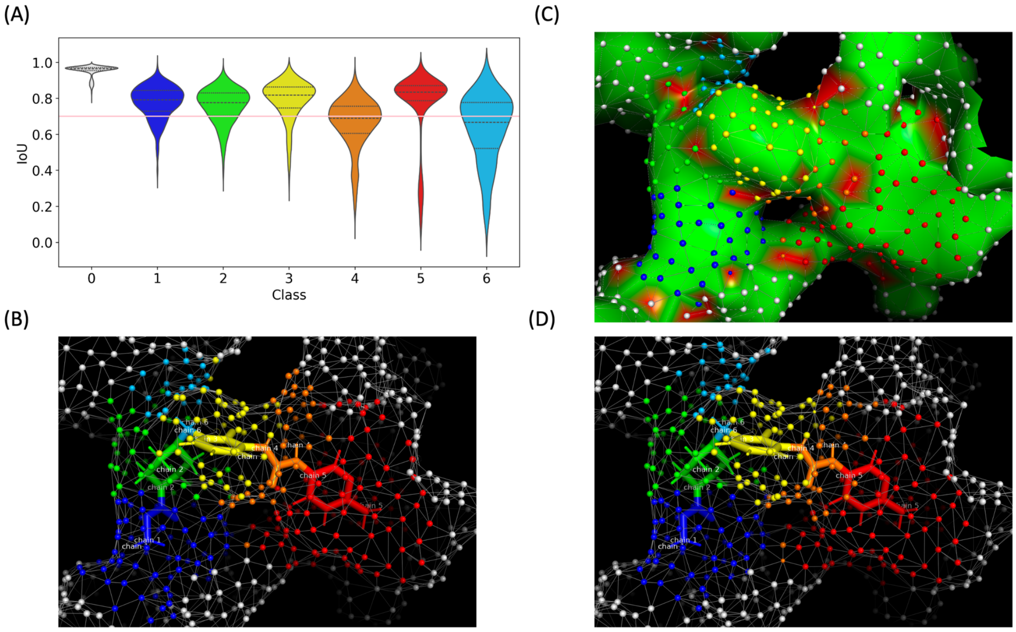


*Figure S3. Performance of the model trained on the holo-PPS-MD1-3 dataset (sampled dataset size: 2400, no 3D random rotation augmentation) on the holo-PPS-MD4 trajectory (test dataset, n = 3066). (A) Distributions of per-class IoU values reported as violin plots (background class, c= 0), a horizontal pink line at 0.7 is included as a visual guide. (B-D) Analysis of model predictions for a sample test frame (holo-PPS-MD4, frame 1410), for which the following performance was observed: mIoU=0.762 (IoU of classes from 0 to 6=[0.963, 0.784, 0.821, 0.741, 0.568, 0.819, 0.636]) and accuracy=94.7%. The molecular surface (SES) in the ROI region is shown as mesh. Vertices are coloured according to the ground truth (B) and predicted (C and D) labels (same colours as panel A). Surface colouring in C indicates correct (green) and incorrect (red) predictions.*

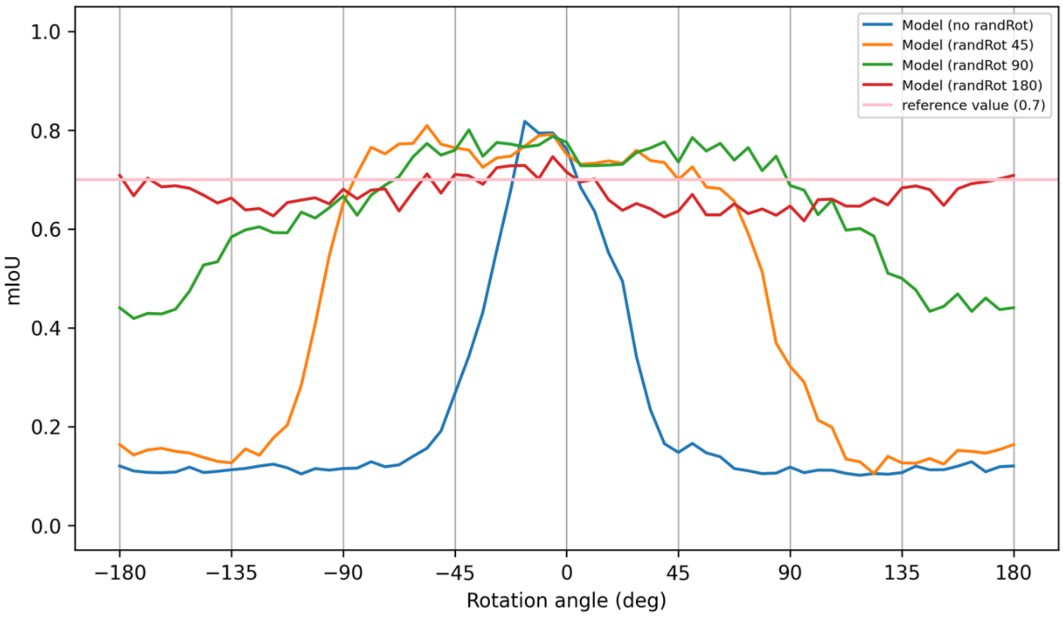


*Figure S4. Performance of the models trained with and without 3D-rotation augmentation when applied to a sample test set frame (holo-PPS-MD4, frame 1410) after rotation around the x axis. Rotation angles sampled from one of four ranges: 0° (no randRot), -45° to 45° (randRot 45), -90° to 90° (randRot 90), or -180° to 180° (randRot 180). A horizontal pink line at 0.7 is included as a visual guide.*

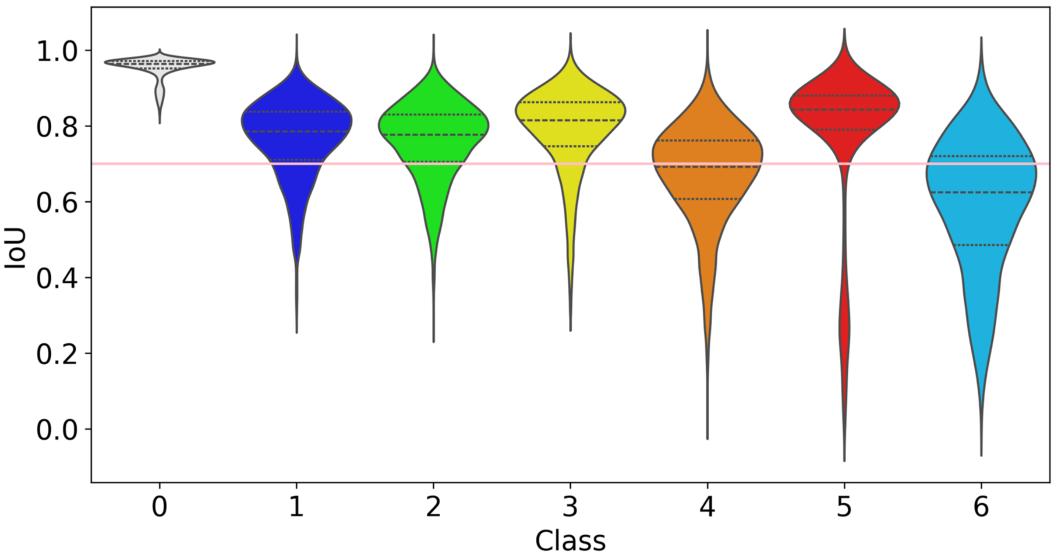


*Figure S5. Performance of the PPS-trained-4800 model on the holo-PPS-MD4 trajectory (test dataset, n = 3066). The model was trained on the holo-PPS-MD1-3 dataset, with a sampled dataset size of 4800, a 3D random rotation range of -180° to 180° and a masked GDF loss. Distributions of per-class IoU values are reported as violin plots (background class, c= 0), a horizontal pink line at 0.7 is included as a visual guide. On this test set, the model achieved a mIoU of 0.756 ± 0.096 and an accuracy of 94.0% ± 3.3%.*

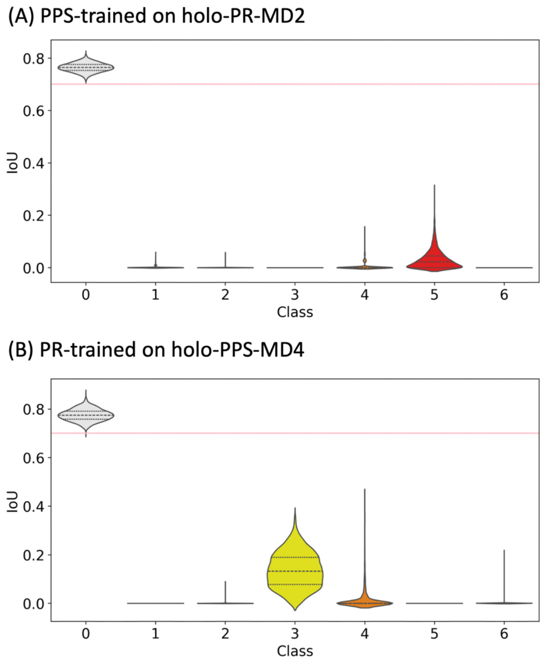


*Figure S6. Performance of single-state models when applied to the opposite state. Violin plots of per-class IoU distributions (n=3066) for the PPS-trained-4800 model applied to the holo-PR-MD2 trajectory (A) (mIoU: 0.114 ± 0.006, accuracy: 73.1 ± 1.3%) and for the PR-trained-4800 model applied to the holo-PPS-MD4 trajectory (B) (mIoU: 0.133 ± 0.013, accuracy: 73.4 ± 1.5%). A horizontal pink line at 0.7 is included as a visual guide.*

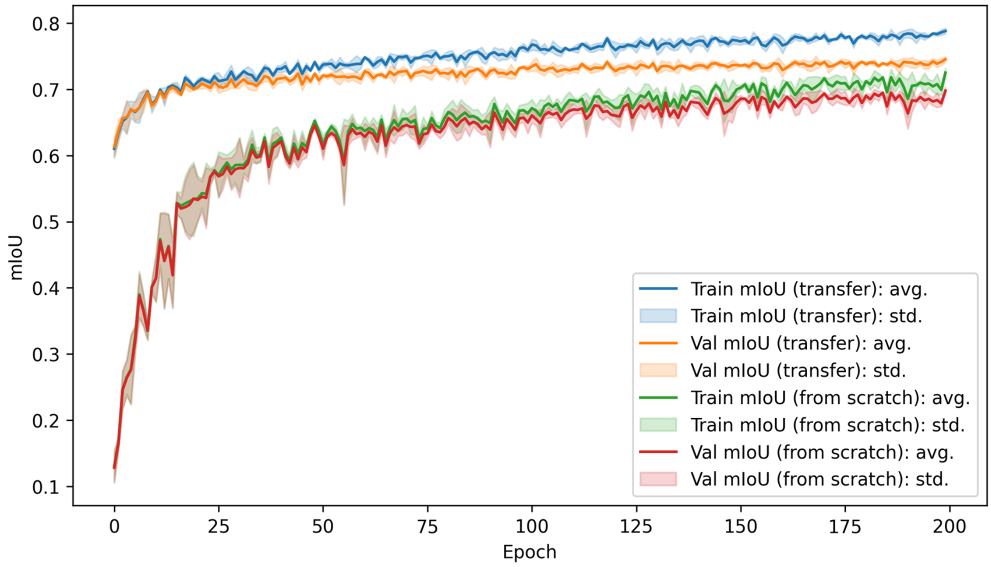


*Figure S7. Training progress over 200 epochs for models trained on the mixed holo-PR-MD1-3 and holo-PPS-MD1-3 dataset (sampled dataset size: 1600, 3-fold cross validation). The training and validation mIoU values averaged over the folds are reported for each epoch, together with their standard deviation (shaded area). The model trained starting from scratch (green and red) is compared to the one retrained from PPS-trained-4800 model weights (blue and orange).*

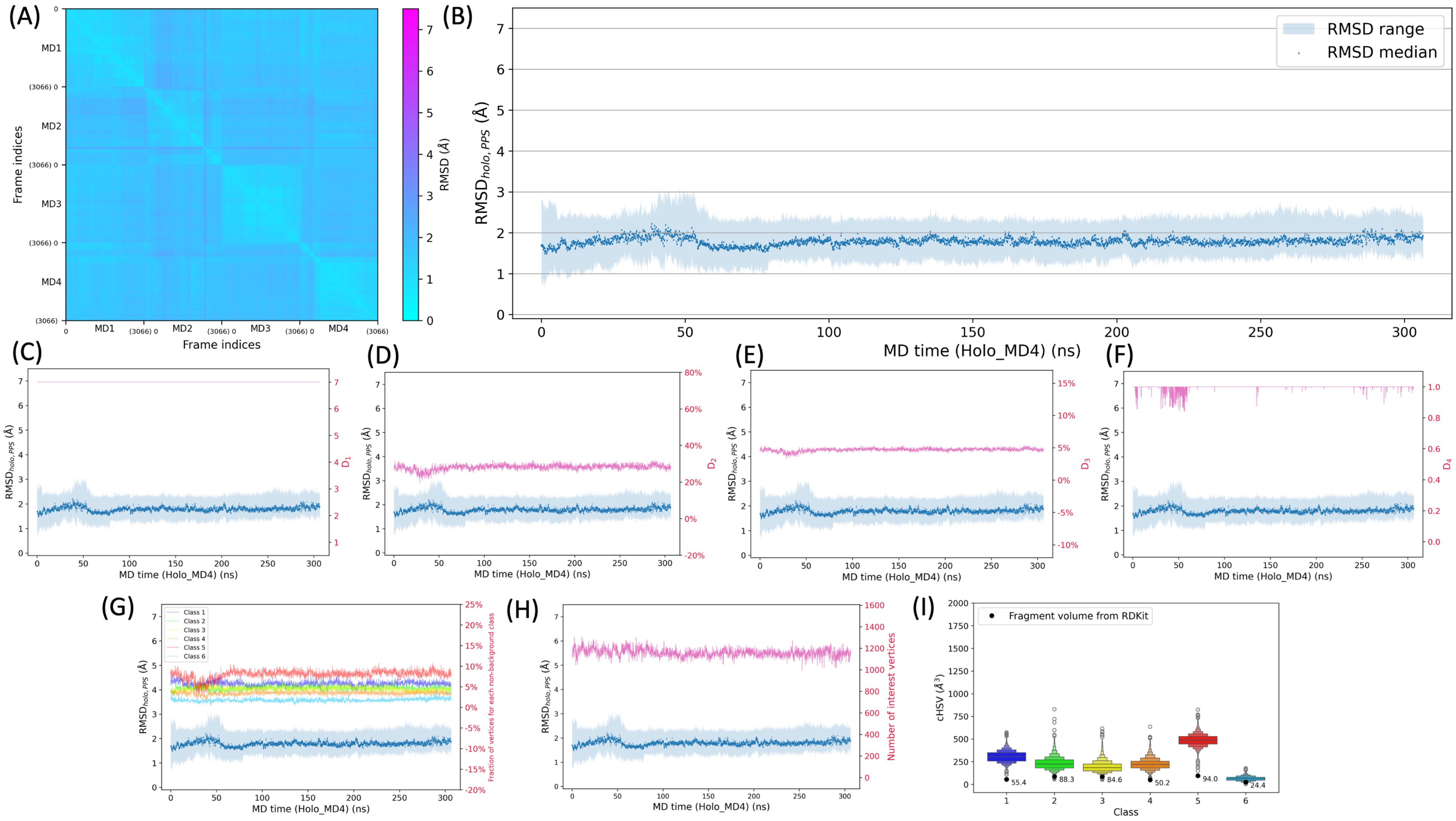


*Figure S8. (A) Pairwise RMSD values (OM binding site only) for the four holo PPS MD trajectories (holo-PPS-MD1-4). (B) Time evolution of $RMSD_{holo,PPS}$ median values (dark blue) during holo-PPS-MD4, with the min-max range indicated as light blue shading. (C-H) Time evolution of deep-learning-derived descriptors (C: $D_1$, D: $D_2$, E: $D_3$, F: $D_4$) and related features (G: fraction of vertices for each non-background class, H: number of ROI vertices) during holo-PPS-MD4. (I) Enhanced box plots of corrected HoloSpace volume ($cHSV_c$) values for each class c compared to the RDKit-calculated volume of the corresponding fragment (black dots) for the holo-PPS-MD4 trajectory.*

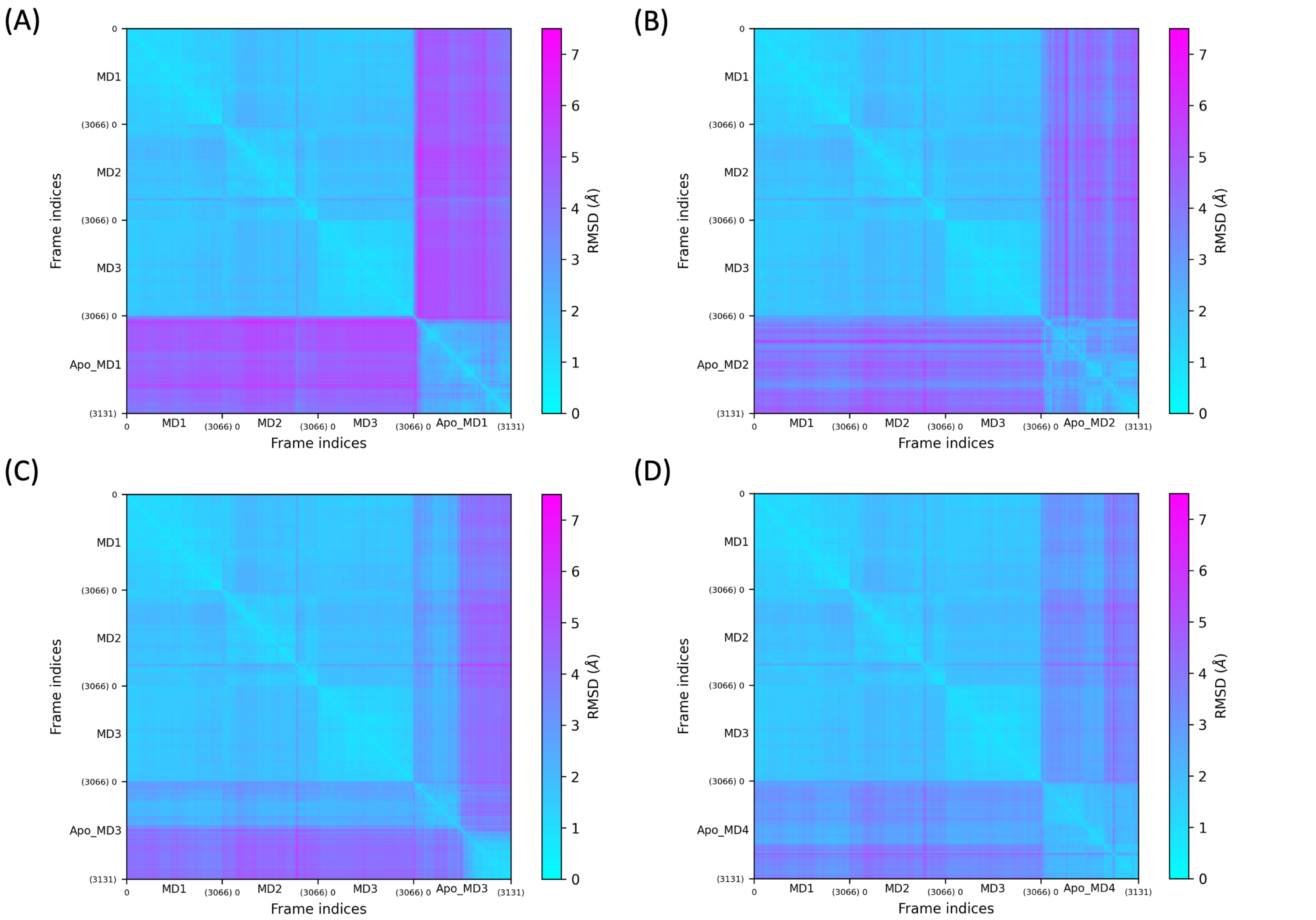


*Figure S9. Pairwise RMSD values (OM binding site only) for three holo PPS trajectories (holo-PPS-MD1-3) and one apo PPS trajectory (A: apo-PPS-MD1, B: apo-PPS-MD2, C: apo-PPS-MD3, D: apo-PPS-MD4).*

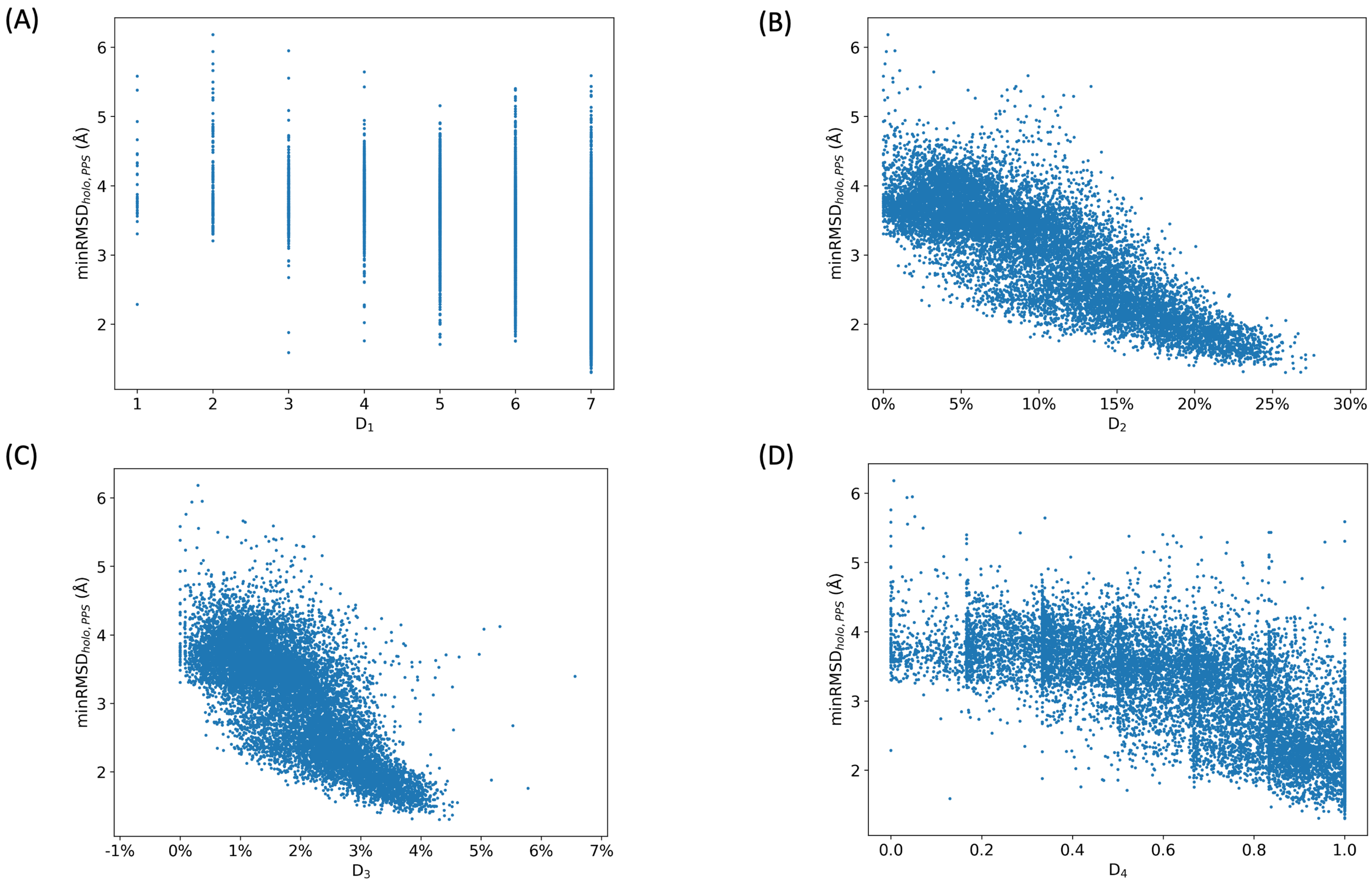


*Figure S10. Scatterplots of minRMSD$_{holo,PPS}$ values against deep-learning-derived descriptors (A: $D_1$, B: $D_2$, C: $D_3$, D: $D_4$) for the apo PPS trajectories (apo-PPS-MD1-4)*

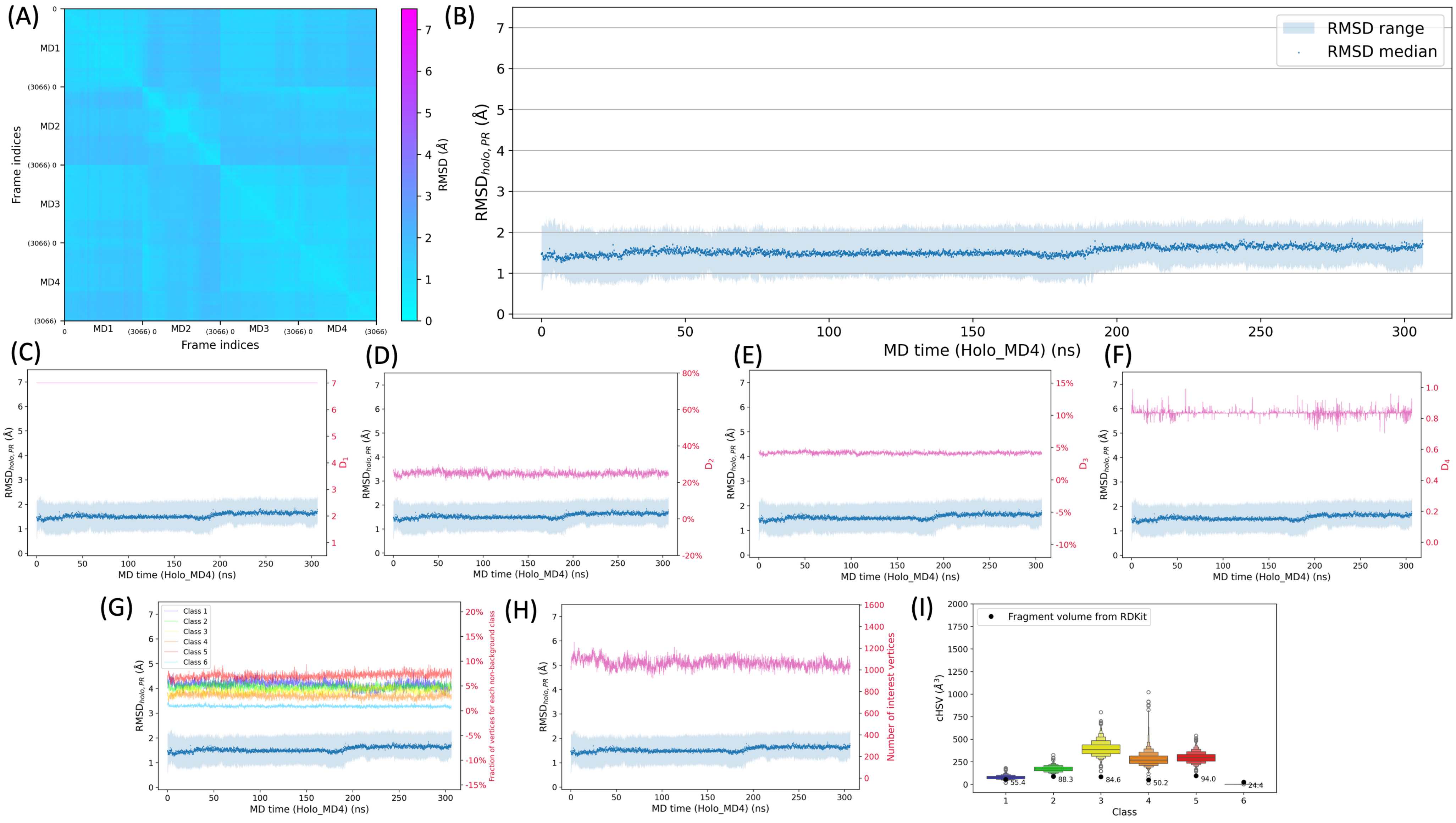


*Figure S11. (A) Raw pairwise RMSD values for the four holo PR MD trajectories (holo-PR-MD1-4). (B) Time evolution of RMSD$_{holo,PR}$ median values (dark blue) during holo-PR-MD4, with the min-max range indicated as light blue shading. (C-H) Time evolution of deep-learning-derived descriptors (C: $D_1$, D: $D_2$, E: $D_3$, F: $D_4$) and related features (G: fraction of vertices for each non-background class, H: number of ROI vertices) during holo-PR-MD4. (I) Enhanced box plots of corrected HoloSpace volume (cHSV$_c$) values for each class c compared to the RDKit-calculated volume of the corresponding fragment (black dots) for the holo-PR-MD4 trajectory.*

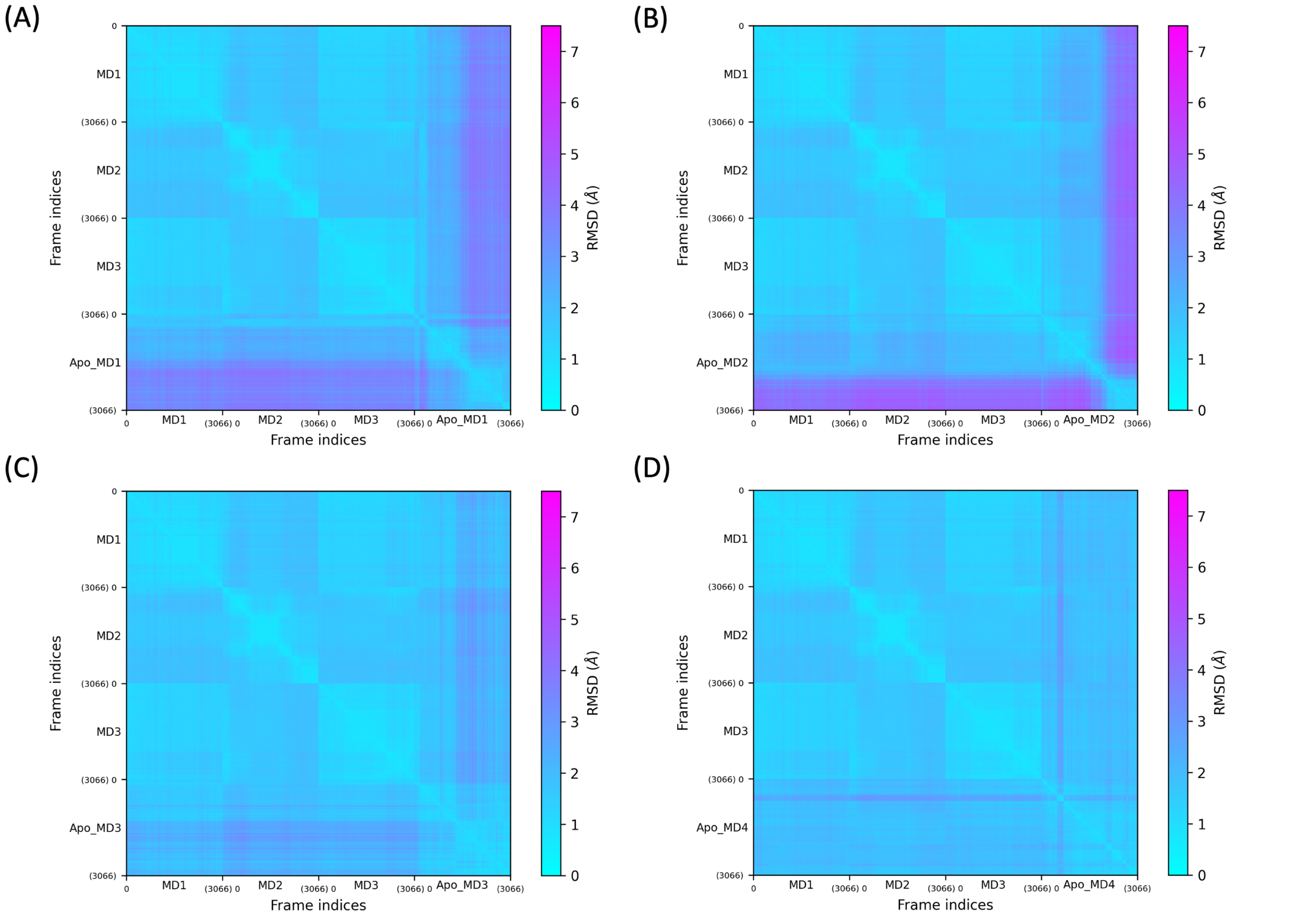


*Figure S12. Pairwise RMSD values (OM binding site only) for three holo PR trajectories (holo-PR-MD1-3) and one apo PR trajectory (A: apo-PR-MD1, B: apo-PR-MD2, C: apo-PR-MD3, D: apo-PR-MD4).*

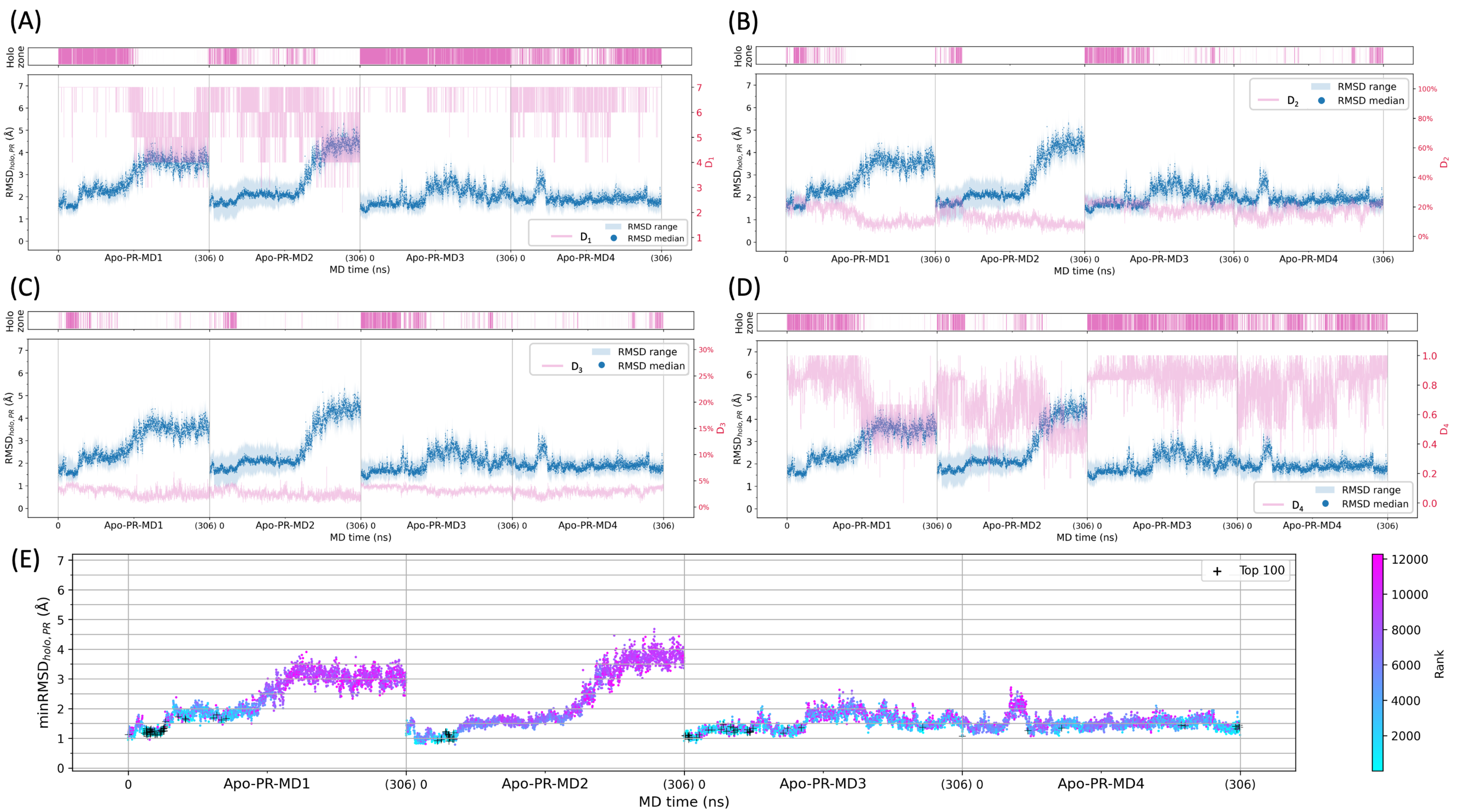


*Figure S13. (A-D) Relationship between RMSD$_{holo,PR}$ values and deep-learning-derived descriptors D$_1$-D$_4$ (pink) for apo-PR-MD1-4 simulations. The time evolution of RMSD$_{holo,PR}$ median values is shown in dark blue, with the min-max range indicated as light blue shading. Dark pink shading in the "holo zone" indicates frames with descriptor values within the holo-PR-MD4 ranges (min-max values in Table S5). (E) Time evolution of minRMSD$_{holo,PR}$ values, where each point is coloured from cyan to magenta according to the R$_{holo}$ rank of the frame. The top 100-ranked frames are indicated with a black cross.*

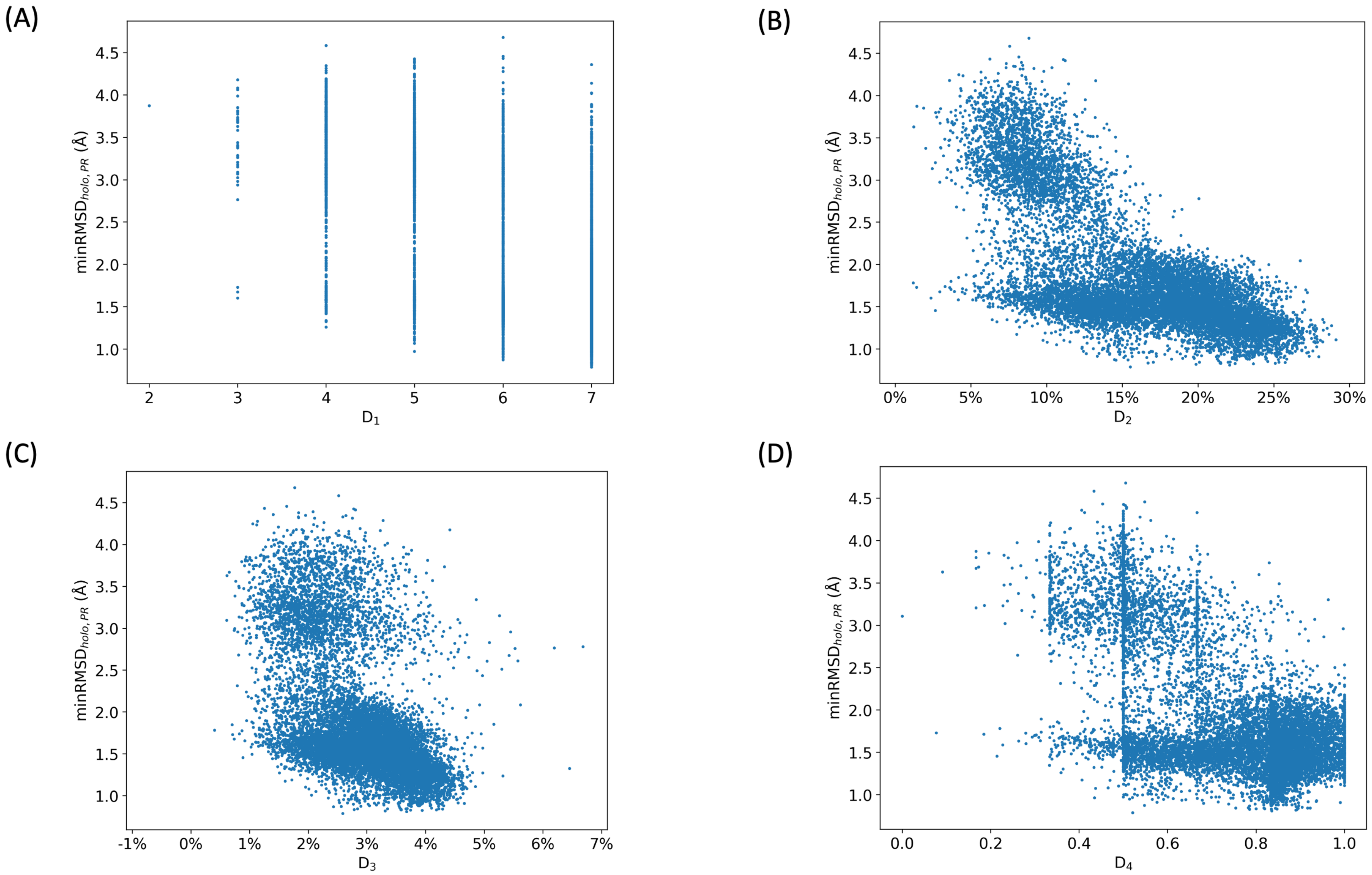


*Figure S14. Scatterplots of minRMSD$_{holo,PR}$ values against deep-learning-derived descriptors (A: $D_1$, B: $D_2$, C: $D_3$, D: $D_4$) for the apo PR trajectories (apo-PR-MD1-4).*

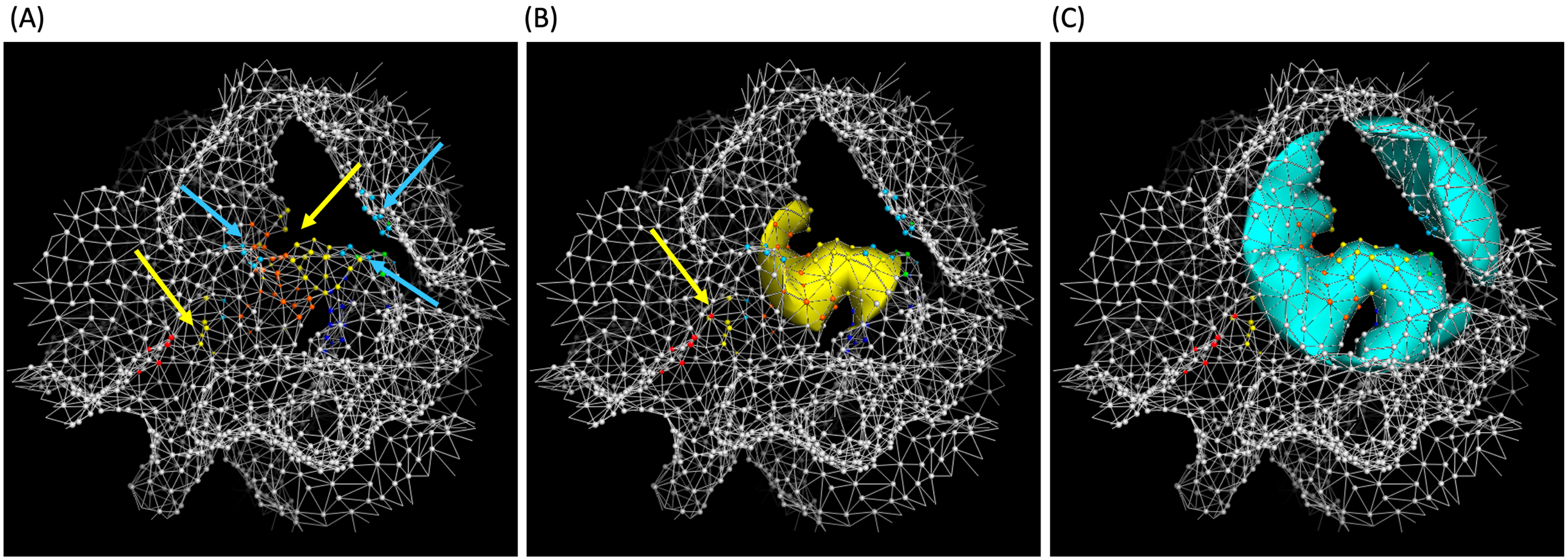


*Figure S15. Example of frame with HoloSpace overestimation (apo-PPS-MD2, frame 2832). The molecular surface (SES) in the ROI region is shown as mesh, with vertices coloured according to the predicted class (see Figure 1B for the colour code, vertices labelled as background are shown in white). (A) Non-contiguous regions labelled as class 3 (yellow) and class 6 (cyan) are highlighted with arrows. (B/C) Solid surfaces are used to represent the class 3 (B) and class 6 (C) raw HoloSpace. In B, class 3 vertices recognised as outliers by DBSCAN are indicated with an arrow.*

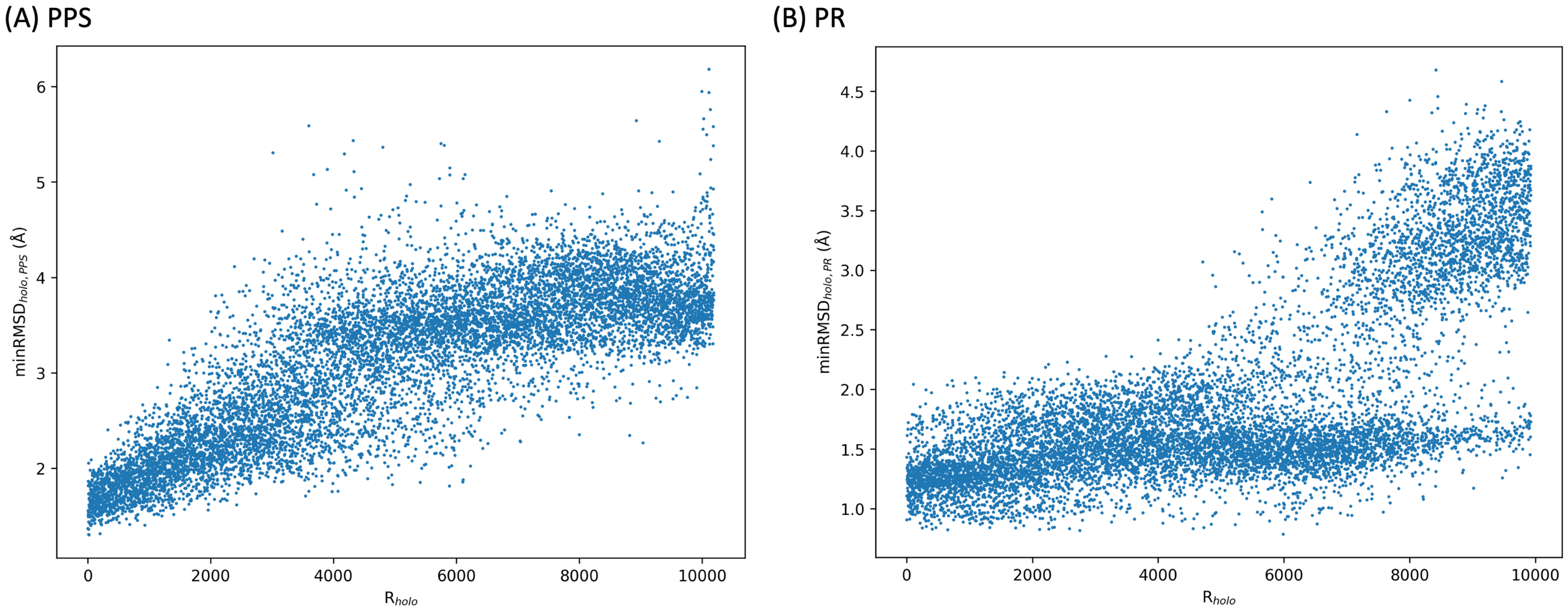


*Figure S16. Scatterplots of minRMSD$_{holo,PPS}$ (A) and minRMSD$_{holo,PR}$ (B) values against $R_{holo}$ rankings for apo-PPS-MD1-4 (A) and apo-PR-MD1-4 (B) trajectories.*

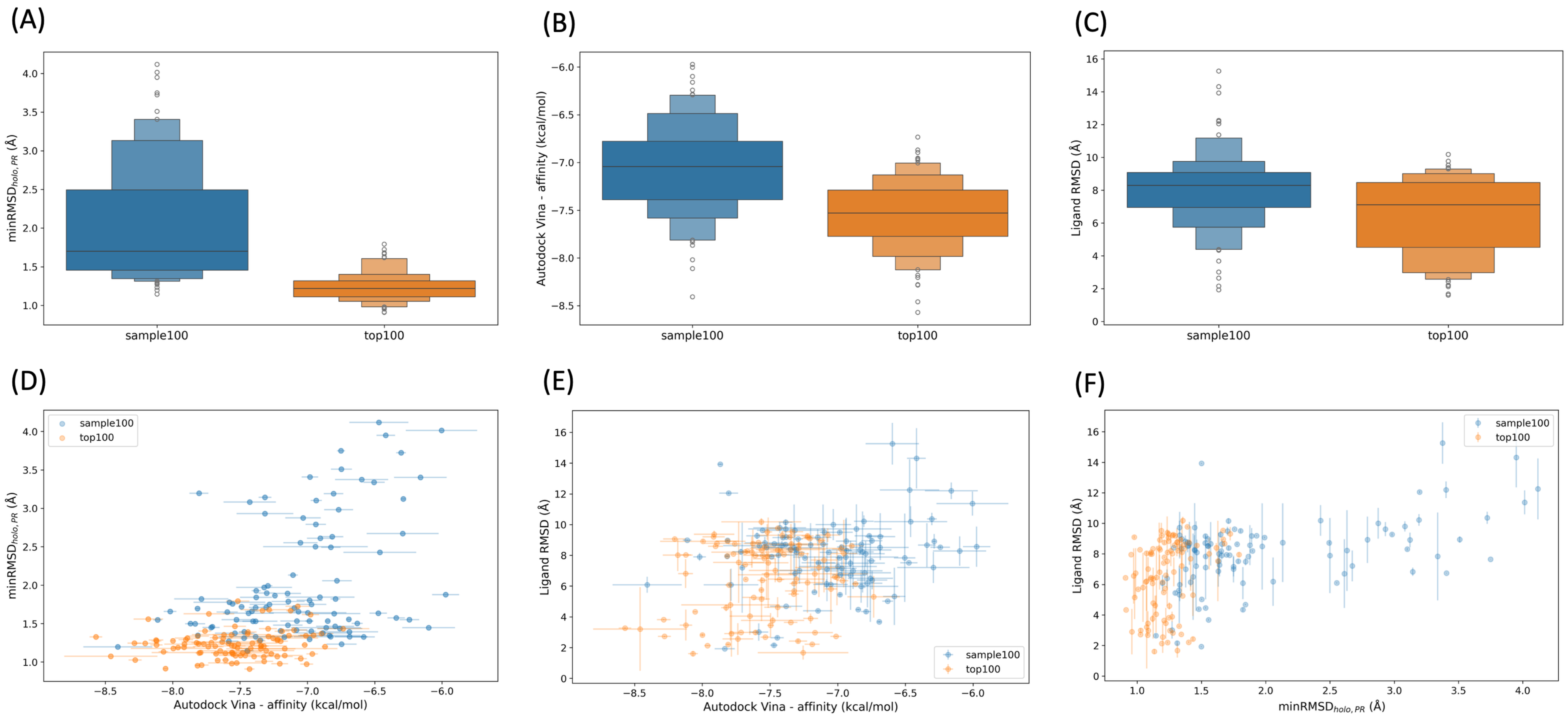


*Figure S17. OM docking to selected frames from apo-PR-MD1-4 trajectories. (A) Enhanced box plots of minRMSD$_{holo,PR}$ values for sample100 (blue) and top100 (orange) frames. (B) Enhanced box plots of Vina binding affinities of OM to sample100 and top100 frames. For each frame, the binding affinity is obtained by averaging the values calculated for the best docking poses from 5 runs. (C) Enhanced box plots of ligand RMSD values relative to the reference holo structure for OM docked to sample100 and top100 frames. RMSD values are calculated using heavy atoms only. (D-F) Scatter plots showing pairwise relationships between minimum RMSD$_{holo,PR}$, Vina binding affinity and ligand RMSD values to the reference holo structure. Error bars show standard deviations across 5 docking runs (using the best docking pose for each run).*

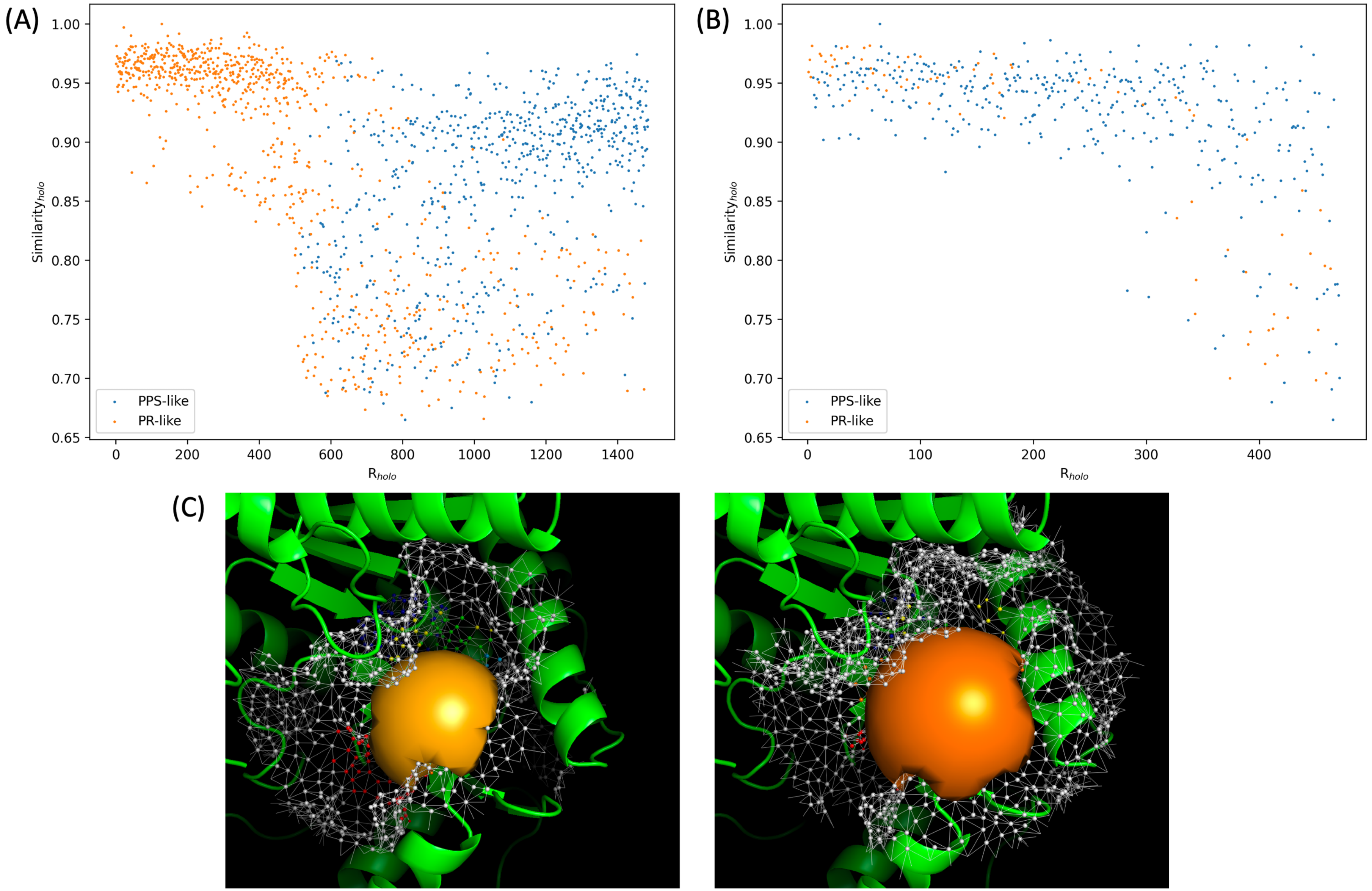


*Figure S18. (A-B) Scatterplot of $Similarity_{holo}$ values against $R_{holo}$ rankings for apo PR-to-PPS trajectory frames. Values obtained using the PR (A) and the PPS (B) ROI definition are plotted. For each plot and frame, the larger of $Similarity_{holo,\ PR}$ and $Similarity_{holo,\ PPS}$ is used. (C) Example of an apo PR-to-PPS frame (frame 230, PR-like) that raises pre-ranking warnings when the PPS ROI definition is used. The raw HoloSpace delimited by class 4 ROI vertices obtained with the PR (left panel) and PPS (right panel) definition is shown as orange surface. The molecular surface (SES) in the ROI region is shown as mesh.*

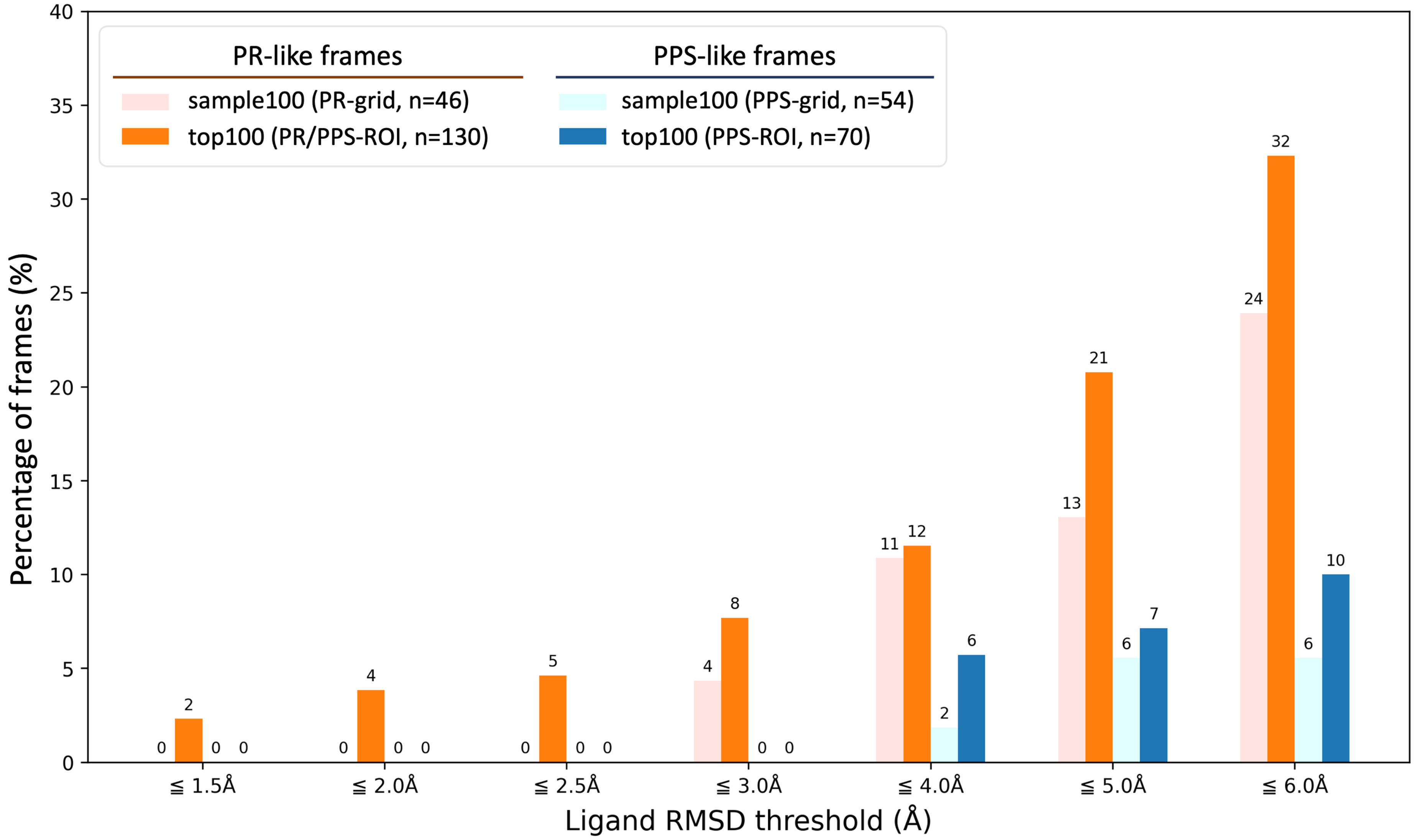


*Figure S19. Percentage of frames with ligand RMSD below a given threshold for sample100 and top100 sets of apo PR-to-PPS frames. Results are grouped by PR-like (orange shades) and PPS-like (blue shades) frames, with control results from the randomly selected set (sample100) shown in the lightest shade. Frames grouped under the top100 PR-like label (dark orange) come from both PR and PPS ROI definitions. It is to be noted that the performance for the 30 PR-like top100 frames from PPS-ROI was likely suboptimal because docking used a grid (PPS-grid) consistent with the ROI definition but not with the actual conformation of the frames.*
